# Extended Nucleic Acid (exNA): A Novel, Biologically Compatible Backbone that Significantly Enhances Oligonucleotide Efficacy *in vivo*

**DOI:** 10.1101/2023.05.26.542506

**Authors:** Ken Yamada, Vignesh Narayan Hariharan, Jillian Caiazzi, Rachael Miller, Chantal Furguson, Ellen Sapp, Hassan Fakih, Qi Tan, Nozomi Yamada, Raymond C. Furgal, Joseph Paquette, Brianna Bramato, Nicholas McHugh, Ashley Summers, Clemens Lochmann, Bruno M.D.C. Godinho, Samuel Hildebrand, Dimas Echeverria, Matthew R. Hassler, Julia F. Alterman, Marian DiFiglia, Neil Aronin, Anastasia Khvorova

## Abstract

Metabolic stabilization of therapeutic oligonucleotides requires both sugar and backbone modifications, where phosphorothioate (PS) is the only backbone chemistry used in the clinic. Here, we describe the discovery, synthesis, and characterization of a novel biologically compatible backbone, extended nucleic acid (exNA). Upon exNA precursor scale up, exNA incorporation is fully compatible with common nucleic acid synthetic protocols. The novel backbone is orthogonal to PS and shows profound stabilization against 3’- and 5’-exonucleases. Using small interfering RNAs (siRNAs) as an example, we show exNA is tolerated at most nucleotide positions and profoundly improves in vivo efficacy. A combined exNA-PS backbone enhances siRNA resistance to serum 3’-exonuclease by ∼32-fold over PS backbone and >1000-fold over the natural phosphodiester backbone, thereby enhancing tissue exposure (∼6-fold), tissues accumulation (4- to 20-fold), and potency both systemically and in brain. The improved potency and durability imparted by exNA opens more tissues and indications to oligonucleotide-driven therapeutic interventions.

## Introduction

Oligonucleotide therapeutics, including small interfering RNAs (siRNAs), are a new class of medicine that enable the modulation of disease-causing genes^1-5^. Five siRNAs have been FDA-approved, with many more candidates in late-stage clinical trials^5-6^. The basis of successful oligonucleotide therapeutic platforms is an optimized chemical architecture that imparts metabolic stabilization to support robust, safe, and sustained modulation of gene expression in a tissue of interest^1,5,7-14^. Phosphorothioate (PS) modification of the oligonucleotide backbone (i.e., internucleotide phosphodiester linkages) is currently the default strategy for stabilization of the 5’- and 3’-ends of oligonucleotides, despite several backbone modifications being discovered. Indeed, PS is the only backbone modification used in clinic^1,5,15^, in part because the PS backbone is structurally similar to the canonical phosphodiester (PO) backbone, enabling high compatibility with RNA-binding protein machinery, such as Argonaute 2 (Ago2), needed for gene silencing^12,15,16-19^. Incorporating a few terminal PS-backbone modifications significantly enhances 5’ and 3’-exonuclease resistance *in vivo*, extending duration of effect from weeks to months^12,19^. However, 3’-truncated metabolites are detectable *in vivo* following months-long duration^18,19^. This finding suggests 3’-end degradation continues to impede long-term efficacy and durability of oligonucleotide therapeutics, perhaps because PS does not alter backbone structure enough to prevent nuclease recognition.

In nature, single carbon (methyl group) attachment to nucleobases and amino acids is used to slightly alter the local structure of DNA, RNA, and protein to modulate nucleic acid-protein interactions and regulate gene expression^20^ (**Fig. 1**). This biological process inspired us to apply an analogous approach to the backbone of siRNA with the goal of reducing interaction with nucleases (to enhance stability) while maintaining interaction with Ago2 (to maintain efficacy).

**Fig 1.**
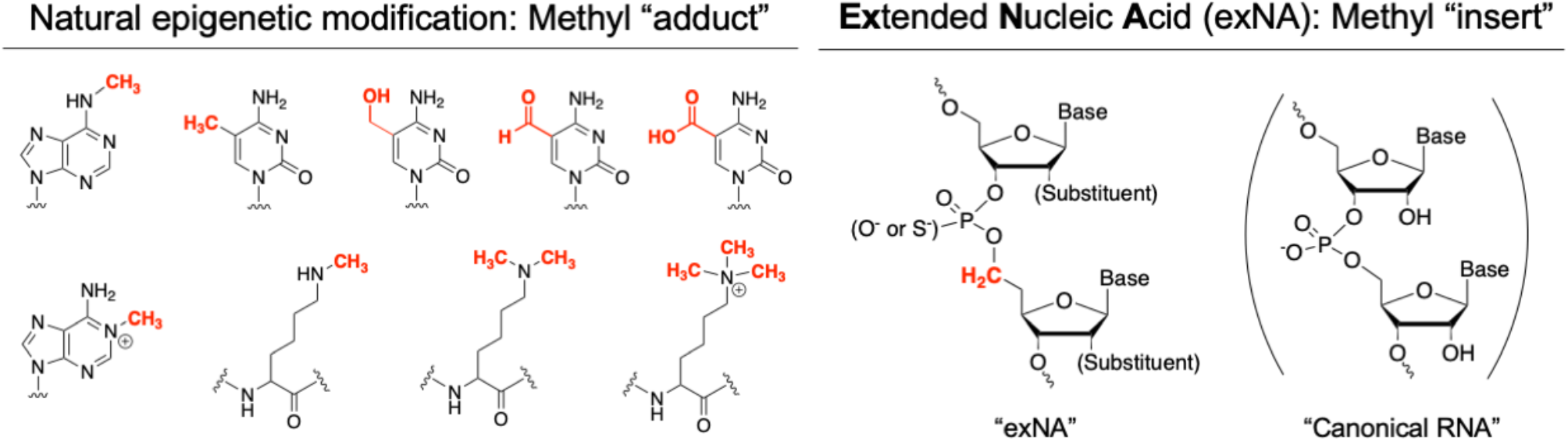
Chemical structure of extended Nucleic Acid (exNA) with methyl inserts (permanent structural modulation) and natural epigenetic modification with methyl adduct (removable by endogenous demethylase).

Here, we describe the discovery, synthesis, and evaluation of a novel, structurally extended backbone chemistry named “*<u>ex</u>tended-<u>N</u>ucleic <u>A</u>cid* (exNA)”, where an extra carbon is incorporated 3’ of phosphate (**Fig. 1**)^21-23^. We demonstrate that exNA imparts profound improvements in exonuclease stability, while minimally impacting overall structure, charge, and thermodynamic stability, of oligonucleotides. exNA is well tolerated at most backbone positions in the context of fully chemically modified siRNAs and is orthogonal to PS. Incorporating both exNA and PS at the 3’-terminal profoundly impacts siRNA clearance and tissue accumulation to improve potency and duration of effect. The exNA-PS backbone opens a new chapter in enhanced stabilization of oligonucleotides, widening the clinical utility and application of the oligonucleotide therapeutics class.

## Results

### Synthesis of extended nucleic acid (exNA) analogues

To evaluate the impact of exNA modification in the context of clinically relevant siRNA scaffolds, backbone-extended nucleosides were synthesized in the context of 2’-O-Methyl (2’-OMe) and 2’-Fluoro (2’-F) modified ribose. 2’-OMe and 2’-F nucleoside modifications enhance metabolic stability, are highly compatible with human Ago2, and are widely used in FDA-approved siRNA drugs and siRNA drug candidates in late-stage clinical trials. 2’-OMe-exNA-uridine and 2’-F-exNA-uridine were initially selected as target compounds for synthesis (**Scheme 1**). Briefly, starting with 5’-O-DMTr protected nucleoside, 3’-OH was protected with TBDMS and TBDPS for 2’-OMe and 2’-F nucleoside, respectively. After deprotection of 5’-O-DMTr, 3’-protected nucleoside was oxidized using IBX reagent to yield 5’-aldehyde compound. Then, separately prepared methyltriphenylphosphonium ylide by potassium tert-butoxide and methyltriphenylphosphonium bromide were reacted with 5’-aldehyde to yield vinyl substituted compound **4a** (or **4b)**^**24**^. Compound **4a** (or **4b**) was reacted with 9-borabicyclo[3.3.1]nonane (9-BBN) and subsequently treated with sodium perborate to produce exNA nucleoside **5a** (or **5b**)^**24**^. After 6’-O-DMTr protection, the 3’ protecting group was deprotected by tetrabutylammonium fluoride, then 3’-O-phosphitylation was performed to yield exNA phosphoramidites **7a** or **8b**). To evaluate the impact of exNA structure in a canonical 2’-OH RNA duplex, ribo-uridine exNA phosphoramidite was also prepared by a similar synthetic scheme (*see supplement*).

**Scheme 1.**
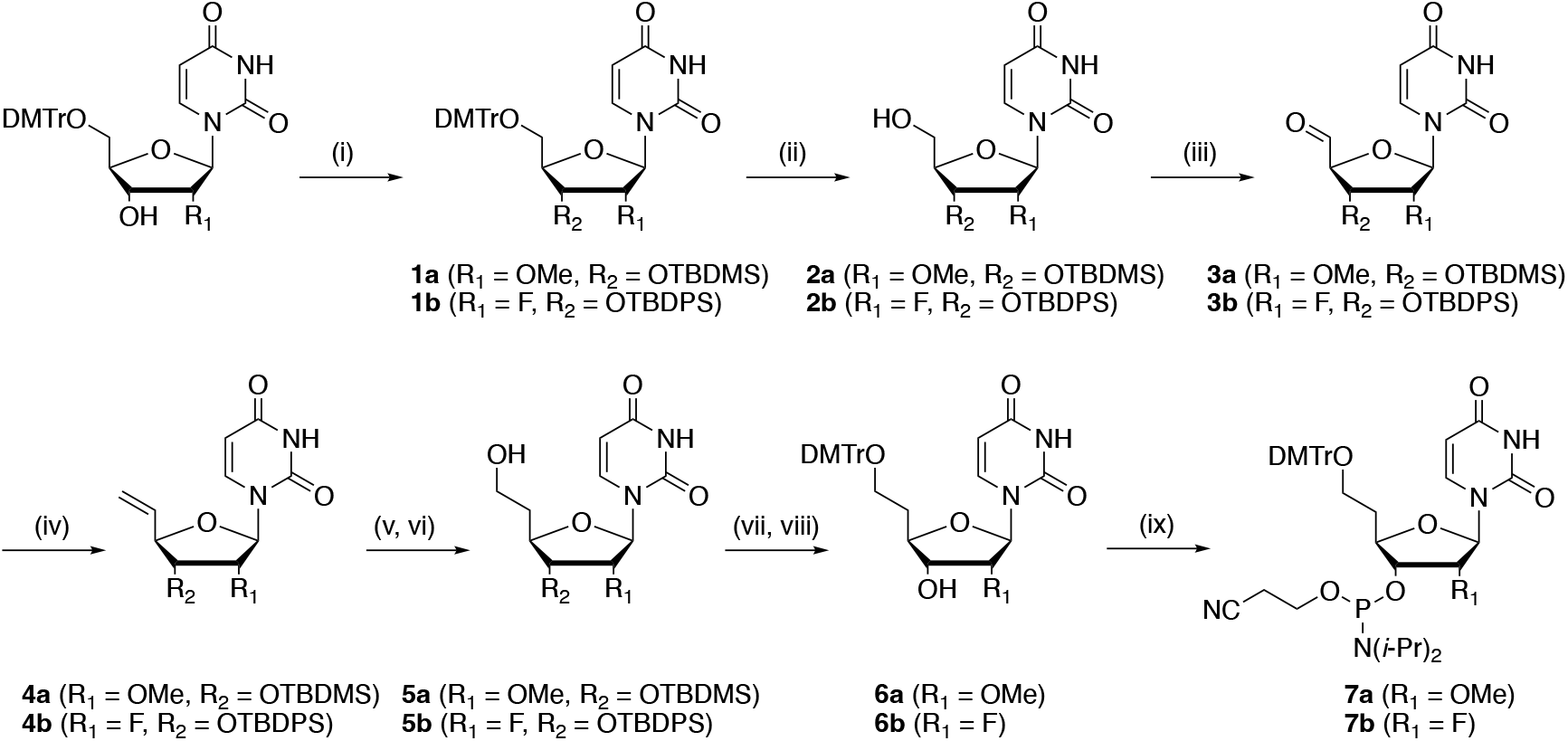
Synthesis of exNA phosphoramidites (**7a** and **7b**). Reagent and conditions: (i) TBDMSCl, Imidazole/DMF, rt, overnight; (ii) 3% DCA/CH_2_Cl_2_, triethylsilane, rt, 1 h, **2a**:88 % (2 steps), **2b**: 84 % (2 steps); (iii) IBX/ CH_3_CN, 85°C, 1.5 h; (iv) CH_3_PPh_3_Br, ^*t*^BuOK, THF, 0°C then rt, overnight, **4a**: 75% (2 steps), **4b**: 67% (2 steps); (v) 9-BBN, THF, 0°C, overnight; (vi) NaBO_3_·4H_2_O, MeOH, THF, H_2_O, 0°C then rt, overnight, **5a**: 62% (2 steps), **5b**: ND; (vii) DMTr-Cl, pyridine, rt, 2 h; (viii) 0.1 M TBAF, THF, rt, 1 h, **6a**:93% (2 steps), **6b:** 12% (3 steps); (ix) 2-cyanoethyl *N,N*-diisopropylchlorophosphoramidite, DIPEA/CH_2_Cl_2_, 0°C then rt, 0.5 h, **7a**: 86%, **7b**: 81%.

### Impact of exNA modification on RNA duplex thermostability

Stability of the siRNA duplex impacts the efficacy of siRNA-mediated gene silencing. Therefore, we sought to understand the general impact of exNA on RNA duplex thermostability. We paired an RNA strand incorporating a single exNA-ribouridine (rxU) with a fully complementary strand or a strand with a single mismatch at the position opposite the exNA insert, and measured thermostability of each duplex. For the fully complementary duplex, rxU had little impact on stability (Δ*T*_m_ = - 3°C), confirming that extra carbon incorporation is well accommodated in the RNA duplex structure. Notably, the destabilizing impact of rxU in the context of a G-U mismatch (wobble) was more pronounced (Δ*T*_m_ = -6°C). exNA incorporation had no effect or reduced the energetic penalty in the other mismatch contexts (Δ*T*_m_ = +5 °C and Δ*T*_m_ = 0 °C for rxU-C and rxU-U, respectively) (**Fig. 2**).

**Fig 2.**
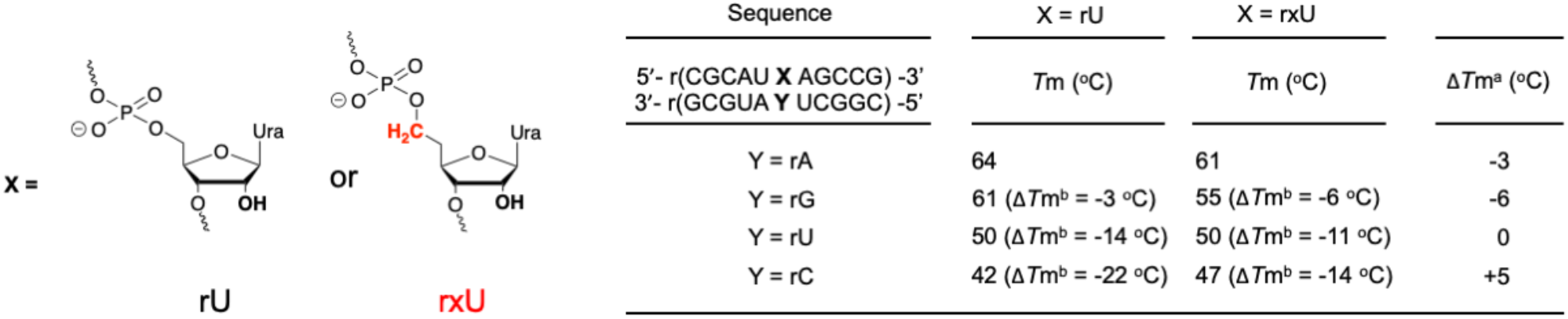
Thermostability of rxU modified RNA duplex. *T*_m_ measurements were carried out in buffer containing 10 mM sodium phosphate (pH 7.0), 100 mM NaCl, 0.1 mM EDTA, and 1 μM duplex. Δ*T*_m_^a^ is the difference in *T*_m_ value between rxU versus rU duplexes. Δ*T*_m_^a^ is the difference in *T*_m_ value between fully complementary duplexes versus duplexes with a mismatch.

### exNA modification profoundly enhances 3’- and 5’-exonuclease resistance in vitro

To evaluate the impact of exNA structure on the exonuclease resistance of oligonucleotides in vitro, we incorporated one or two 2’-OMe modified exNA at the 3’- or 5’-terminus of a model oligonucleotide strand with PO (^ex^PO) or PS (^ex^PS) backbones. Terminal PO and PS modifications without exNA were used in control compounds (see methods). We first tested the stability of each compound against the 3’-exonuclease, snake venom phosphodiesterase I (SVPD). At the conditions tested, the PO control was fully degraded, whereas the PS control showed better stability (Half-life 0.03 vs 1.1 for 3’-PO and 3’-PS, respectively, **Fig. 3A**), consistent with previous reports^25^. Surprisingly, ^ex^PO supported higher 3’-exonuclease resistance than the PS control (∼9 versus 1.1 half-life). Furthermore, ^ex^PS exhibited a half-life that was ∼36-fold higher than the PS control, and ∼1000-fold higher than the PO control. We next tested the stability of each compound against the 5’-exonuclease, bovine spleen phosphodiesterase II (BSP). PO control degraded rapidly, whereas ^ex^PO significantly enhanced 5’-exonuclease resistance but was not as much as PS backbone. (**Fig 3B**) when exNA is incorporated at 1^st^ nucleotide. When exNA incorporated at 2^nd^ nucleotide, the stabilization effect was more prominent than introduction to 1^st^ position. We observed identical stabilization with PS under the condition tested, which is expected as introduction of exNA at 2^nd^ position directly modulate the structure of the first 5’-exonuclease cleavage site.

**Fig 3.**
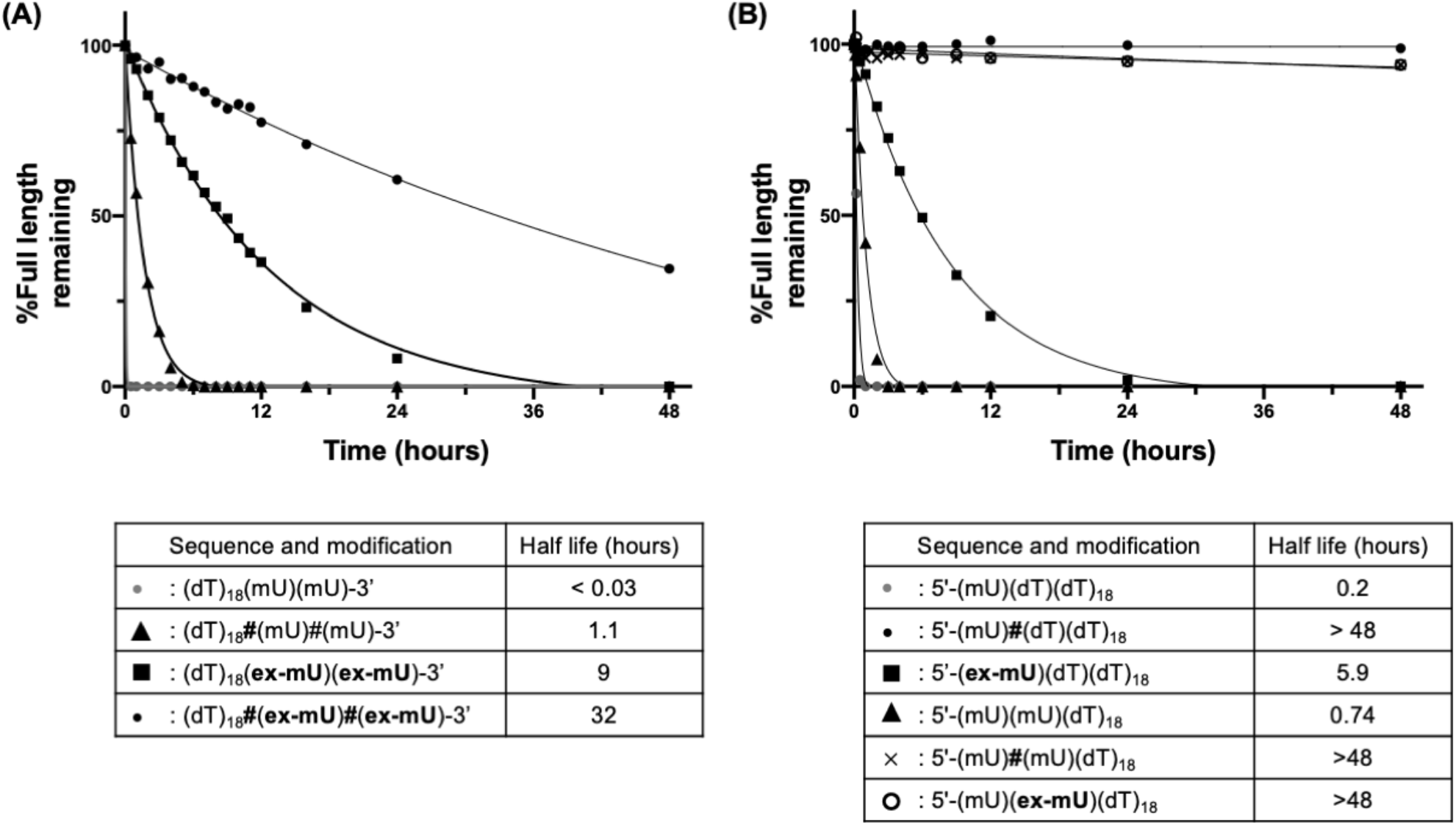
Nuclease resistance of exNA modified oligonucleotides. Stability of oligonucleotide in the presence of (**A**) Snake venom phosphodiesterase I (SVPD) and (B) bovine spleen phosphodiesterase II (BSP). Residual oligonucleotide length was monitored over time and quantified by HPLC. (dT): 2’-deoxy-thymidine, (mU): 2’-OMe-uridine, (**ex-mU**): 2’-OMe-exNA-uridine. “**#**” indicates phosphorothioate linkage, otherwise phosphodiester linkage.

Our results suggest that exNA modification could be used as an alternative to PS for 3’ and 5’stabilization of oligonucleotides or used in combination with PS to impart potentially the highest stability of any biologically compatible backbone reported to date. We elected to further investigate the latter strategy.

### Position-dependent impact of exNA on siRNA efficacy in vitro

During the process of siRNA loading into Ago2 and subsequent target RNA binding and cleavage, each nucleotide of the siRNA has its own specific interaction with the proximal amino acids of Ago2^7, 16, 26-29^. To examine the impact of the extended backbone structure of exNA on siRNA efficacy in cells, we systematically replaced each nucleotide position of a fully chemically modified antisense strand (^**ex**^**AS1**-**20)** or sense strand (^**ex**^**SS1**-**15)** with either 2’-OMe-exNA-Ur or 2’-F-exNA-Ur to create a panel of exNA-containing siRNAs. Moreover, to evaluate the impact of multiple exNA insertions at the 3’-end of the antisense strand, ^**ex**^**AS21**-**24** were designed to contain one, two, three, or four 3’-end exNA inserts, respectively (**Table S4**). All siRNAs contained 2’-OMe or 2’-F ribose modifications and terminal PS linkages (**Fig. 4**) and targeted a previously validated human *Huntingtin* (*Htt*) mRNA sequence (for all sequences, see **Table S4)**^30-32^.

**Fig. 4.**
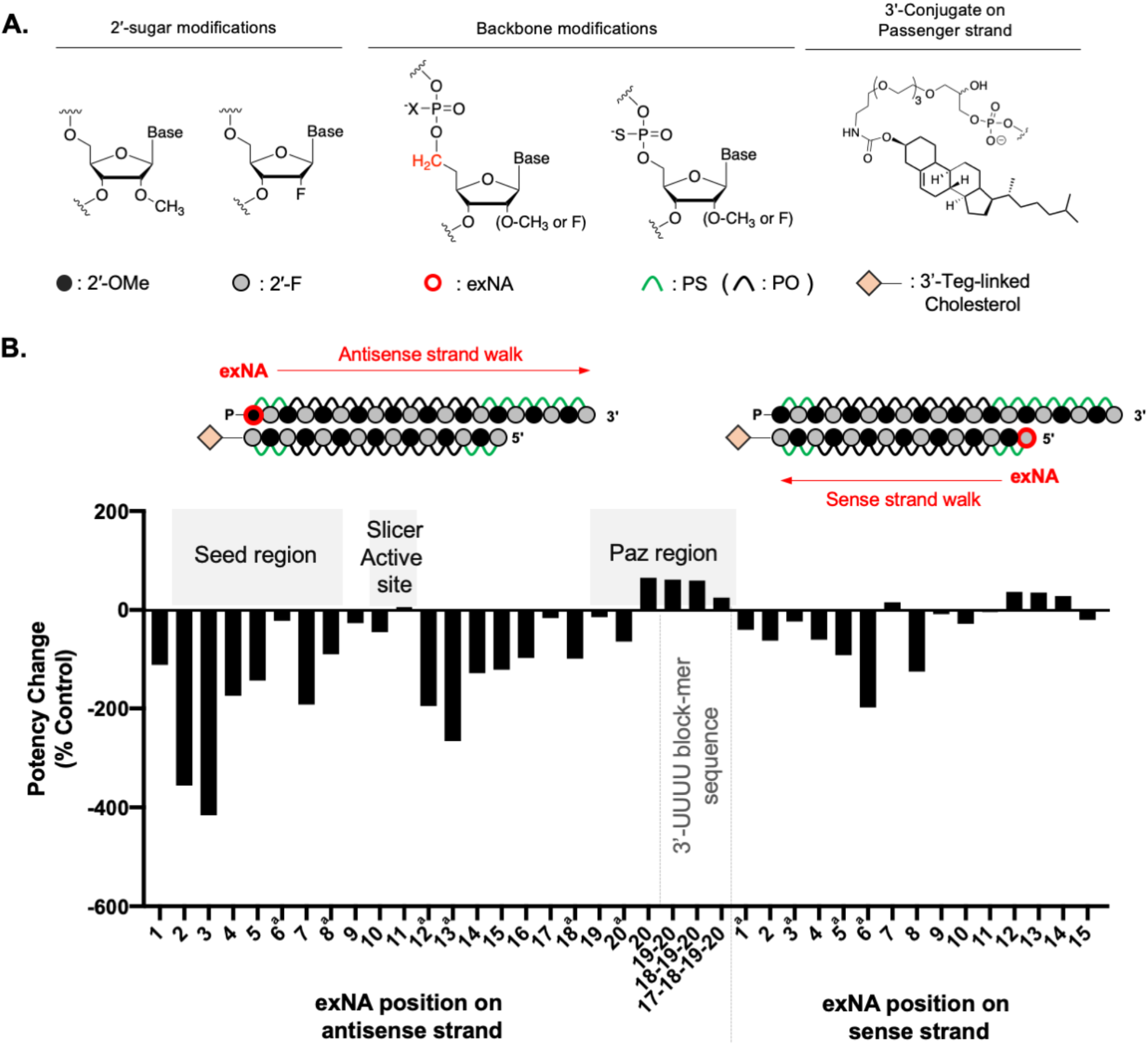
Position-dependent impact of exNA replacement on siRNA activity in vitro. (**A**) Schematic structure of chemical modification used in siRNA. (**B**) Position dependent impact of exNA on siRNA efficacy. Silencing efficacy change was defined by 100 x [IC_50_(Ctrl)-IC_50_(exNA)/IC_50_ (Ctrl)]. A value of ∼0 or a positive value (colored in red) indicate exNA modification was fully compatible with Ago2 or enhanced efficacy, respectively, compared to corresponding control siRNAs. See **Tables S4** and **S5** for sequences, and **Table S6** for all IC_50_ values used to calculate efficacy changes in the figure. ^a^Potency change calculated based on IC_50_ by lipid-mediated uptake.

**Fig. 4** shows the positional impact of exNA incorporation on chemically modified siRNA efficacy, calculated as the difference in IC50 values (derived from a 7-point dose response) for exNA containing siRNA versus the corresponding non-exNA control compound (See methods). exNA was well tolerated in most positions of the sense strand (except position 5) and the antisense strand, except for the seed region (position 2-7) and position 14.

Placement of the extended backbone in the seed region likely reduced silencing efficacy due to interference with base stacking of the seed during target recognition and binding by Ago2, thereby reducing target affinity^16,26,33-36^. The intolerance to structural change at position 14 is harder to rationalize but is consistent with previous work showing this position is sensitive to bulky sugar modifications^13, 37-39^.

Interestingly, the exNA was fully tolerated proximal to the mRNA cleavage site of Ago2 (antisense position 9-10). This finding is consistent with our previous report evaluating another backbone modification, structurally locked internucleotide (*E*)-vinylphosphonate (^i^*E*-VP)^36^. This, combined with the lack of clear crystal structure resolution in this region, suggests that Ago2 can flexibly accommodate both constraining and lengthening backbones at position 9-10^19^. The most exciting observation was the complete tolerance for incorporation of up to four exNA at the 3’ end (**Fig. 4 and Fig. S3**). We even observed a slight but reproducible (with different sequences, data not shown) increase in potency (∼2-fold) with a single or double exNA incorporation. Terminal nucleotides do not contribute to target recognition^40-42^ and the lack of full complementarity at the 3’ terminal caused by exNA may protect the antisense strand from target RNA-induced Ago2 unloading^46^. It is also possible that 3’-end exNA modification may induce better structural fitting into the Paz region of the Ago2 or lower affinity enhance the rate of catalytic cleavage.

### exNA modification block enhances stabilization of fully chemically modified antisense strand against 3’ exonuclease

Typically, stabilization imparted by a chemical modification can be further enhanced when the modification is incorporated in a block (i.e., multiple positions in a row)^43^. Because up to four exNA modifications were tolerated at the 3’-end of the antisense strand in the context of chemically modified siRNA, we sought to better understand the differential impact of 3’-end single, double, triple, and quadruple exNA blocks on antisense strand stability. We analyzed the 3’ stability of ^**ex**^**AS21-24** from our panel of fully modified siRNAs (**Table S4**) using the SVPD assay. We found that exNA incorporation effects were cumulative, where incorporation of two exNA provides further stabilization compared to a single exNA. Further improvement in stability for triple and quadruple exNA incorporation, however, was minimal (**Fig. S4**).

Collective results from in-vitro siRNA efficacy and 3’-exonuclease resistance studies demonstrate that 3’-terminal exNA block modification significantly improves 3’-exonuclease resistance without compromising Ago2 function, suggesting it might enhance overall efficacy and durability of oligonucleotides in vivo. Because enhanced stability seemed to plateau with double exNA incorporation, this block was selected for in vivo evaluation.

### 3’-exNA modification robustly improves the plasma clearance profile of DCA-conjugated siRNA in mice

Upon delivery into the bloodstream, fully modified siRNAs are predominantly degraded by serum 3’-exonuclease^44,45^. Clearance kinetics from the bloodstream profoundly impact the rate of tissue distribution, where higher tissue exposure (Area Under the Curve, AUC) correlates with higher tissue accumulation. We therefore sought to evaluate the effect of 3’-exNA modification on oligonucleotide clearance kinetics in the bloodstream in vivo.

The previously characterized *Htt*-targeting fully chemically modified siRNAs^30^ were synthesized with and without a double 3’-exNA-PS block (**Table S8, Fig. 5A**) and conjugated to docosanoic acid (DCA), which enables widespread tissue distribution to, and efficacy in, extrahepatic tissues^46^. A 10 mg/kg dose of DCA conjugated fully modified siRNAs without double exNA (**D49, or PS**) or with double exNA (**D50, or exNA-PS**) was subcutaneously administered to FVB mice (see methods), and plasma samples were collected at 0, 5, 15, and 30 minutes, and 1, 2, 3, 6, 9, and 24 hours (**Fig. 5B**). Incorporation of exNA had a profound impact on siRNA plasma clearance profile; **D50** exhibited a ∼3-fold enhancement in C_max_ (504 vs 1660 pmol/mL), 6.3-fold increase in AUC_0-inf_ (164 vs 1034 pmol/mL·min), and 2-fold shift (187 vs 305 min) in mean residence time compared to **D49** (**Fig. 5C**). Our findings confirm the significant, negative impact of 3’ degradation during siRNA absorption and distribution, and demonstrate the effectiveness of exNA in inhibiting this degradation.

**Fig. 5.**
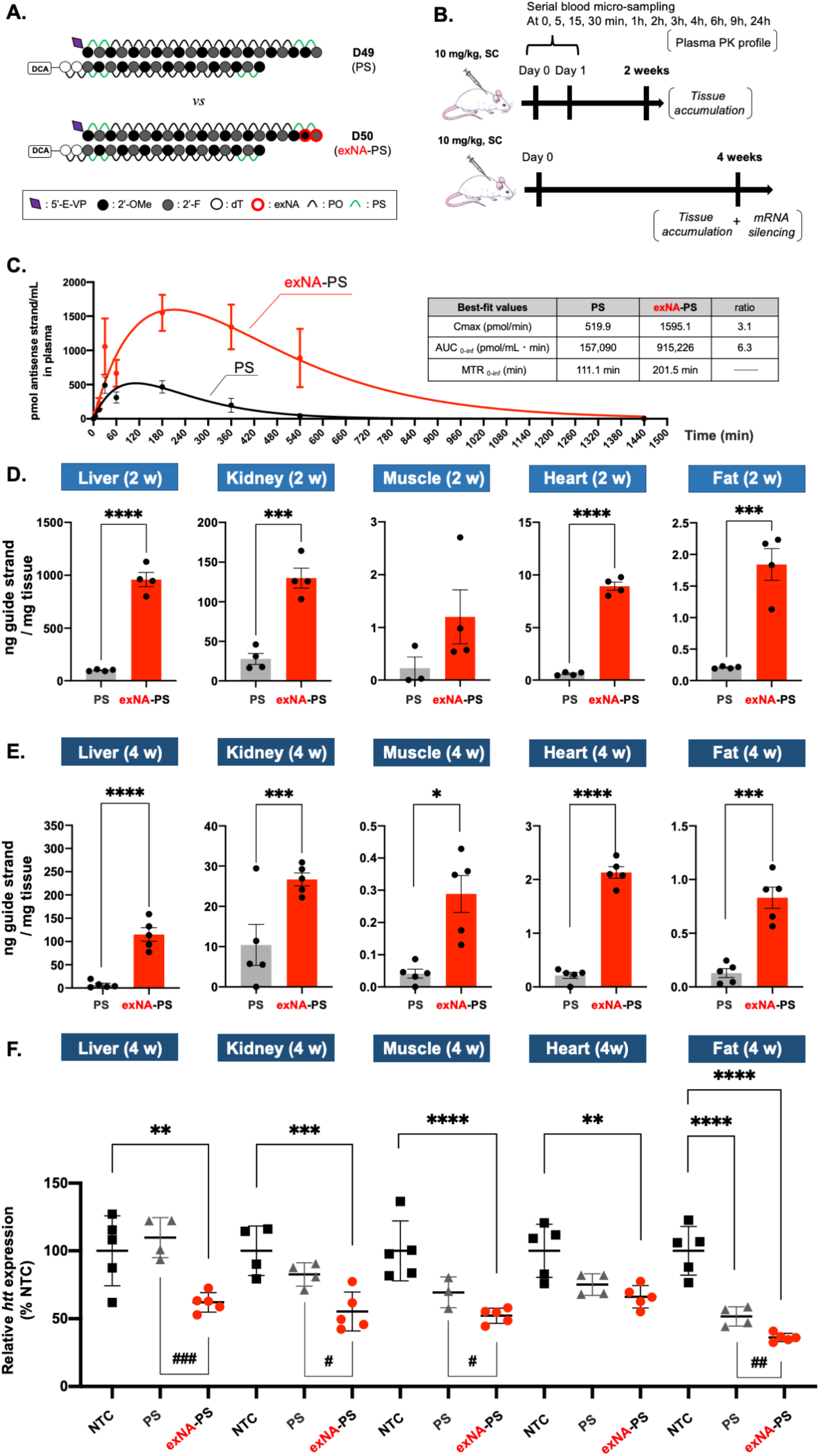
Systemic delivery of DCA-conjugated siRNA with 3’-exNA modification. FVB mice injected with 10mg/kg of DCA conjugated siRNA variants subcutaneously. (A) Schematic of the siRNA scaffolds used, **(B)** Graphical study outline. (**C, D and E**) siRNA antisense strand accumulation in (C) Plasma concentrations of antisense strand (left) and tabulated curve fitting parameters (right) at 5min, 15min, 30min, 1h, 3h, 6h, 9h and 24h post injection, (**D**) tissues at 2 weeks and (**E**) tissues at4 weeks post injection. (**F**) *Htt* mRNA level at 4 weeks post injection measured by Quantigene 2.0 assay. NTC: Non-targeting control. n=4 (C and D) and n=5 (E and F). Data represented as mean ± s.d. of individual animals. Statistical significance calculated for individual tissues using two-tailed unpaired t-test (D and E) or ordinary one-way ANOVA with multiple comparisons (F). Pairwise comparisons between PS and exNA-PS in (F) were performed using two-tailed unpaired t-test (*/# -p<0.05, **/## -p<0.01, ***/### -p<0.001, ****/#### -p<0.0001).

### 3’-exNA modification robustly enhances DCA-conjugated siRNA accumulation levels in multiple tissues

To determine whether the profound effect of exNA modification on plasma clearance kinetics translated into changes in tissue accumulation, mice used for measuring plasma pharmacokinetics were sacrificed at 2 weeks post-injection and siRNA accumulation in tissues was evaluated by the PNA hybridization assay^47^. **D50** (exNA-PS) exhibited 9.7-, 4.7-, 3.5-, 14.9-, and 8.9-fold higher accumulation in liver, kidney, muscle, heart, and fat, respectively, compared to **D49** (PS) (**Fig. 5D**). To confirm whether enhanced tissue accumulation of **D50** is maintained over a longer time course, **D49** and **D50** were administered subcutaneously (10 mg/kg) and tissues were collected at 4 weeks post-injection. **D50** exhibited 16-, 2-, 5.8-, 7.9-, and 6.4-fold higher accumulation than **D49** in liver, kidney, heart, muscle, and fat, respectively (**Fig. 5E**). Our results indicate that simple addition of two carbons (double exNA modification) to the 3’ end of siRNA chemical architecture can profoundly enhance liver and extrahepatic tissue accumulation and retention.

### 3’-exNA modification robustly enhances the potency of DCA-conjugated siRNA in mice

To determine whether the observed enhancement in tissue accumulation by 3’-exNA modification translated into functional modulation of target gene expression, we also quantified *Htt* mRNA expression in collected tissues at 4 weeks post injection (10 mg/kg of D49 or D50). We intentionally selected a relatively low dose and longer timeline for gene expression analysis to increase the likelihood of detecting potential differences in **D49** and **D50** silencing efficacy. At the selected dose and timeline, **D49** did not induce statistically significant silencing (compared to non-targeting control) in any tissue, except for fat. By contrast, **D50** supported productive silencing in all tissues tested (45%, 48%, 34% and 64% in liver, kidney, muscle, heart and fat respectively), with statistically significant difference observed between **D49** and **D50** in all tissues but heart (**Fig. 5F**).

To confirm that the effects are transferable to other targets, we synthesized previously-validated fully chemically modified (including PS) siRNA targeting *myostatin* without and with 3’-exNA (**D52** and **D53**, respectively, **Table S8**) and found that exNA modification significantly enhanced myostatin silencing in quadriceps (**Fig. S5**). Thus, the effects of this new backbone stabilization on extrahepatic gene silencing efficacy are reproducible in a different target and sequence context.

### 3’-exNA modification enhances siRNA potency in mouse central nervous system (CNS)

To evaluate the generalizability of the observed efficacy improvement in the context of a different siRNA scaffold, we synthesized Apolipoprotein E (*ApoE*) and *Htt* targeting siRNA in a di-valent, CNS-active configuration discovered by our group^48^. In this di-valent scaffold, two siRNAs are covalently connected through a linker at the 3’ end of the sense strand. The increase in size and cooperativity of cellular interactions enables broad distribution in the CNS and potent target silencing. Fully chemically modified di-valent siRNA targeting *ApoE* and *Htt* were synthesized with and without 3’-end double exNA modification (exNA-PS or PS, respectively, **Fig. 6A, Table S8**).

**Figure 6.**
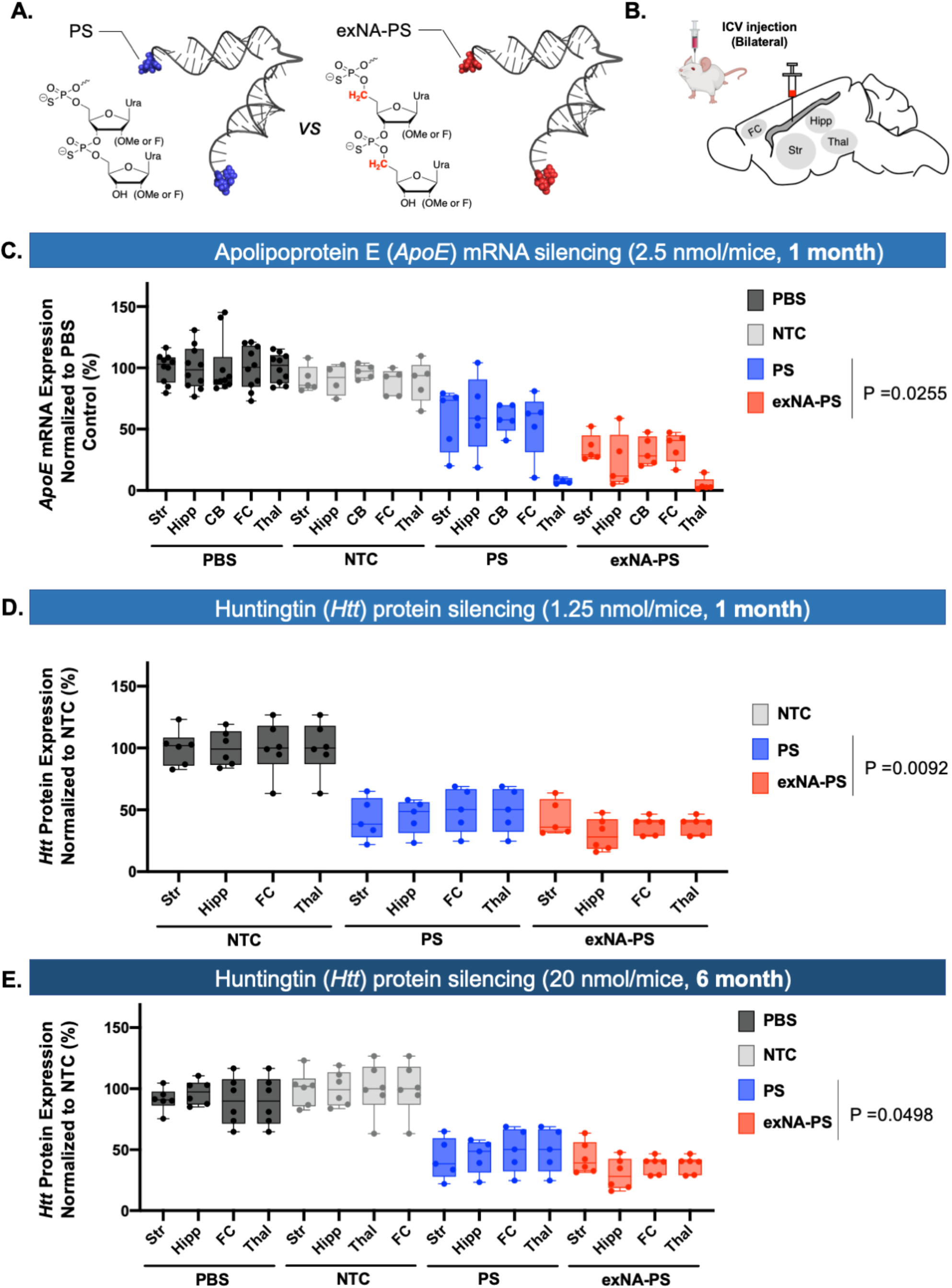
Impact of 3’-exNA modification on siRNA potency in the CNS. (**A**) Schematic of di-valent siRNA structure without and with 3’-exNA modification. (**B**) Brain region of interest. FC: Frontal cortex; Str: Striatum; Thal: Thalamus; Hipp: Hippocampus. (**C**) WT mice injected with 2.5 nmol/10 μL of PS (**D55**) or exNA-PS siRNAs (**D56**) targeting ApoE at 8 weeks of age. Levels of ApoE mRNA were evaluated using Quantigene at 1 month post injection. glmmTMB results: log(exNA effect)=-0.663, P=0.0255. (**D**) WT mice injected with 1.25nmol/10 μL of PS (**D58**) or exNA-PS siRNAs (**D59**) targeting Htt at 8 weeks of age. The levels of WT huntingtin protein in frontal cortex, striatum, thalamus, and hippocampus were evaluated using western blot (ProteinSimple) 1-month post injection. N = 5, one-way Anova with Tukey correction for multiple comparisons. glmmTMB results: log(exNA effect)=-0.519, P=0.0092. (**E**) YAC128-HD mice^49^ injected with 20nmol/10μL with PS or exNA-PS siRNAs targeting Htt at 8 weeks of age. The levels of WT huntingtin protein in frontal cortex, striatum, thalamus, and hippocampus were evaluated using western blot (ProteinSimple) six months following siRNA injections. N = 6, one-way Anova with Tukey correction for multiple comparisons. glmmTMB results: log(exNA effect)=-0.321, P=0.0498. For sequences of di-siRNAs used, see **Table S8**.

Fully chemically modified di-valent siRNA targeting *ApoE* and *Htt* were previously shown to induce robust silencing at a 5-nmol dose. To increase the likelihood of detecting differences in silencing efficacy between PS and exNA-PS divalent siRNAs, we delivered compounds at lower concentrations (2.5 nmol and 1.25 nmol for *ApoE*-targeting and *Htt*-targeting compounds, respectively) to the CNS via intracerebroventricular injection (**Fig 6B**). Htt-RNAs exist both in nucleus and cytoplasm, both of which can be detected by Quantigene 2.0 assay. Thus, we measured *Htt* protein expression level, which directly associates with cytoplasmic *Htt*-mRNA silencing level. At one-month post injection, gene silencing was observed in all regions (except thalamus), and silencing efficacy was enhanced by the addition of 3’ terminal exNA. The effects and trends were similar for both *ApoE-* and *Htt*-targeting compounds (**Fig. 6C-D**), confirming that the effects of exNA modification are sequence and target independent. To confirm whether enhancement of efficacy by exNA modification is maintained over a longer time frame, we injected 20 nmols of exNA-PS or PS siRNAs (*Htt*-targeting) into mice, and measured *Htt* expression at six-months post injection. Consistent with one-month results, silencing efficacy was significantly enhanced by the addition of 3’ terminal exNA (**Fig. 6E**).

Collectively, our results suggest that 3’ terminal exNA modification enhances siRNA efficacy, potency, and durability independent of target sequence and scaffold.

## Discussion

Metabolic stability is critical to the clinical success of oligonucleotide drugs. Although many stability-enhancing backbone modifications have been identified, including phosphoroamidate morpholino oligonucleotides (PMOs), Peptide Nucleic Acids (PNAs), Phosphoramidate (PN), and Mesyl-phosphoramidates (PN-MS), PS is largely the only siRNA backbone modification used in the clinic. This is primarily because PS provides significant stabilization and modulates PK properties^50^ without impacting PO-backbone charge and architecture (e.g., spacing between phosphates), enabling siRNA to maintain compatibility with Ago2. However, 5’ and 3’ exonucleases recognize nucleic acid backbone by sensing the relative distance between phosphate charges^51-53^, and PS modified oligonucleotides appear to be susceptible to 3’ nucleases after months-long duration in vivo.

To investigate the strategy to further enhance metabolic stability without compromising siRNA efficacy, we describe here a novel non-natural nucleic acid backbone, exNA, where an extra carbon is inserted between the 5’-OH and 5’-carbon of the nucleoside. This extra carbon insertion has no impact on charge and does not introduce significant stereo-constraint, preventing disruption to Watson-crick base pairing. As a result, exNA has minimal impact on duplex thermostability, enhances tolerance for mismatches, and is compatible with Ago2. Indeed, exNA shows compatibility with Ago2 in a wider range of backbone positions in the siRNA compared to a previously published inter-nucleotide modification, (*E*)-vinylphosphonate (^i^*E*-VP), suggesting an extended flexible backbone structure is better tolerated than a structurally constrained backbone. On the other hand, the extra carbon in exNA slightly extends the relative distance between backbone phosphate charges, causing a more profound stabilizing effect against 3’-exonuclease than that of PS and equal stabilization effect against 5’-exonuclease. Thus, exNA represents a new class of backbone chemistry with potential application in engineering therapeutic oligonucleotides.

Whereas exNA may inhibit initial nuclease binding to the oligonucleotide, PS induces metabolic stabilization by destabilizing the formation of reaction intermediates when the nuclease cleaves the backbone. Our results, therefore, suggest that blockage of initial nuclease binding has a more profound impact on the nuclease resistance properties of oligonucleotides than that of the stabilizing mechanism of PS. However, exNA is an orthogonal modification to PS, presenting an opportunity to combine exNA and PS in one linkage to inhibit nucleases by two different mechanisms. Indeed, we find that backbones with combined exNA-PS modification synergistically enhances 3’-exonuclease resistance. The enhanced stability effect seems to plateau with two exNA-PS modifications per strand, which could suggest that the rate-limiting step of 3’-exonuclease is binding to the oligonucleotide. Indeed, the co-crystal structure of 3’-exonuclease with substrate shows that only the first few nucleotides are recognized ^51^. Our work suggests than an exNA-PS combinatorial backbone may represent the best option for extensive oligonucleotide stability.

Although our current evaluation focused on exNA and exNA-PS backbone modifications in the context of siRNAs, the susceptibility to degradation by 5’ and 3’ nucleases is shared across the oligonucleotide therapeutics class. Thus, the enhanced stability and pharmacokinetics/dynamics (PK/PD) conferred by terminal exNA-PS modification should be transferable to CRISPR guides, ASOs, mRNAs, tRNAs, and other RNA therapies^54-56^. Interactions between oligonucleotides and their effector proteins (Cas9, RNaseH, etc.) are less transferable across the therapeutic class. Therefore, future work must systematically investigate the tolerability of exNA incorporation at internal positions in other RNA therapies.

There are numerous chemistries being explored for oligonucleotide therapies. However, the clinical utility of these chemistries depends on manufacturability. exNA modification is orthogonal to not only PS, 2’-OMe, and 2’-F, but also 2’-methoxyethyl (MOE), bridged (or locked) nucleic acids (BNAs and LNAs), PN, Ms-PN backbones, and other reported chemical modifications^5,57^. Here, we focus on most commonly used 2’-OMe, 2’-F, and PS in clinic, but further characterization of other combinations is expected to produce a similar level of impact on plasma PK profiles, tissue accumulation, and potencies. Another important factor dictating utility is ease/scalability of synthesis. exNA nucleosides can be synthesized from commercially available nucleosides in a straightforward manner. Indeed, we demonstrate that exNA conversion can be done in just 2 steps (Wittig reaction and Hydroboration). exNA phosphoramidites require no specific condition to incorporate into oligonucleotides on solid support, which can be handled the same way as commercially available nucleoside phosphoramidites.

One of our most striking observations was the impact of 3’ exNA modification on plasma clearance kinetics, which resulted in significantly enhanced tissue accumulation at both 2- and 4-weeks post injection. This result is likely due to enhanced stability against 3’ exonuclease and indicates that 3’ degradation during absorption and distribution significantly impacts tissue accumulation of oligonucleotides. However, it is also possible that exNA-PS modification alters plasma protein binding properties to impact PK properties and tissue accumulation, which should be investigated in future work.

Oligonucleotide delivery and functionality are well established for liver, with the tri-*N*-acetyl-galactosamine (tri-GalNAc) conjugate serving as the basis for several FDA approved and clinically advanced siRNA drugs^5^. Therefore, demands in the RNA therapeutics field have shifted towards delivery to extrahepatic tissues. Here, we show that, even with non-optimal conjugate mediated delivery, the enhanced tissue accumulation of exNA-PS modified siRNAs translates into improved functional efficacy in extrahepatic tissues. This observation was not dependent on delivery configuration, as we see it for systemic delivery with a hydrophobic conjugate and direct administration into cerebrospinal fluid with a di-valent siRNA scaffold; and thus, is likely applicable to other delivery modalities (e.g., antibodies, peptides, etc.). The enhanced efficacy and durability of paired exNA and PS modification may therefore offer an opportunity to advance the clinical utility of RNA drugs in extrahepatic tissues.

In summary, exNA is a synthetically accessible, scalable, and biologically compatible backbone that provides profound stability against nucleases. This modification has the potential to expand the clinical utility of siRNA and potentially other oligonucleotide therapies to more tissues and indications.

## ACKNOWLEDGEMENTS

This project was funded by the NIH/NINDS [R01 NS104022 for A.K.]; S10 [OD020012 for A.K.]; MIRA [R35 GM131839 for A.K.]; NIH CREATE [U01 NS114098 for N.A.]; CHDI Foundation [RecID A-5038 for N.A.], the Berman–Topper Fund (for A.K. and N.A.). We would like to thank the University of Massachusetts Chan Medical School Animal Medicine Department and veterinary technicians for their contributions to the large-animal studies. We would like to thank Dr. Emily Haberlin for editing the manuscript, Dr Akbar Ali for maintaining infrastructure for NMR, and Dr. Yanglan Tan and Dr. SonNgoc Nguyen for measurement of high-resolution mass analysis.

## Author contributions

K.Y. and A.K. conceived the project. K.Y., V.N.H., and A.K. contributed experimental design for exNA studies in hepatic and extrahepatic tissues, and K.Y. V.N.H. J.C., H.F., Q.T., R.C.F, A.S., C.L, B.M.D.C.G. conducted the experiments. K.Y., R.M., C.F., E.S., J.F.A., M.D., N.A. and A.K. contributed experimental design for exNA studies in CNS, and R.M. and E.S. conducted experiments. K.Y. and N.Y. contributed to exNA thermal stability and nuclease resistance study. K.Y. N.Y.,B.B., N.M., and D.E. contributed oligonucleotide synthesis. S.H. contributed statistics analysis of exNA CNS studies. K.Y. and A.K. wrote the manuscript.

## Methods

### Oligonucleotide synthesis

All oligonucleotides were synthesized on a MerMade 12 (BioAutomation) synthesizer on 1-10 μmol scale using universal UNY Linker support or cholesterol 3’-lcaa CPG (ChemGenes). Synthesis was conducted on a standard 1, 5, or 10 μmol scale RNA phosphoramidite synthesis cycle, which consists of (i) detritylation, (ii) coupling, (iii) capping, and (iv) iodine oxidation to phosphate by 0.02 M I_2_ in THF-pyridine-H_2_O (7:2:1, v/v/v) or sulfurization by 0.1 M DDTT in pyridine-acetonitrile (9:1, v/v). For monomer building blocks, commercially available 2’-OMe or 2’-F-modified nucleoside phosphoramidites and custom synthesized 2’-OMe or 2’-F exNA-phosphoramidites were used. For antisense strands used in the in-vitro screen, 5’-terminus was coupled with phosphorylating reagent, bis(2-cyanoethyl)-N,N-diisopropylphosphoramidite (ChemGenes) to have 5’-phosphate. Metabolic stability of 5’-terminal phosphate against endogenous phosphatase is an essential parameter that defines potency and duration of effect of siRNA in vivo. 5’-(*E*)-vinylphosphonate (VP) is a well-known phosphatase-resistant 5’-phosphate analogue allowing for prolonged duration of effect of siRNA in vivo. Thus, for all guide strands used in in-vivo experiments, 5’-vinyl tetra phosphonate (pivaloyloxymethyl) and 2’-*O*-methyl uridine 3’-CE phosphoramidite (Hongene Biotechnology Co., Ltd.) were coupled as a 5’-terminal nucleotide to generate the 5’-(*E*)-VP moiety. The coupling of all monomers building blocks were conducted under standard conditions on solid support using 5-(benzylthio)-1-*H*-tetrazole (BTT) as an activator. Cholesterol-conjugated oligonucleotides were prepared on a 500-Å LCAA-CPG support, where the cholesterol moiety is bound to tetra-ethylenglycol through a succinate linker (ChemGenes, co.). For DCA conjugated oligonucleotide synthesis, docosanoic acid was directly attached via an amide bond to a controlled pore glass (CPG) functionalized by a C7 linker, as described^58^. For divalent sense strand synthesis, in-house synthesized solid support was used as described^48^. For all 2’-OH RNA synthesis, standard 2’-TBDMS protected phosphoramidites and exNA-Ur phosphoramidite were used and coupled by a standard RNA synthesis and deprotection protocol^59^. Synthesized 5’-phosphate guide strands and 3’-cholesterol conjugated passenger strands were deprotected by AMA (conc. NH_4_OH -40% MeNH_2_ in water, 1:1, v/v) at 26°C for 2 h. Guide strand bearing 5’-VP was conducted in conc. NH_4_OH (3% diethylamine) at 35°C for 20 h. Crude guide strands and cholesterol-conjugated passenger strands were purified by anion-exchange HPLC and reversed-phase HPLC, respectively.

### LC–MS analysis of oligonucleotides

Identities of oligonucleotides were authenticated by LC–MS analysis on an Agilent 6530 accurate mass Q-TOF LC–MS machine using the following conditions: buffer A: 100 mM hexafluoroisopropanol/9 mM triethylamine in LC–MS grade water; buffer B:100 mM hexafluoroisopropanol/9 mM triethylamine in LC–MS grade methanol; column, Agilent AdvanceBio oligonucleotides C18; gradient antisense, 0% B 1 min, 0–30% B 8 min, clean and re-equilibration 4 min; gradient sense, 0% B 1 min, 0–50% B 0.5 min, 50– 100% B 8 min, clean and re-equilibration 4 min; temperature, 45 °C; flow rate, 0.5 ml min−1, UV (260 nm). MS parameters: source, electrospray ionization; ion polarity, negative mode; range, 100–3,200 *m*/*z*; scan rate, 2 spectra s−1; capillary voltage, 4,000; fragmentor, 180V. All reagents were purchased from Sigma Aldrich and used per manufacturer’s instructions unless otherwise stated.

### Thermostability assay

1 μM guide strand and 1 μM sense strand were annealed in a 10 mM sodium phosphate buffer (pH 7.2) containing 100 mM NaCl and 0.1 mM EDTA by heating at 95°C for 1 min and cooled down gradually to room temperature. *T*_m_ measurement was performed with a temperature controller. Both heating and cooling curves were measured over a temperature range from 20 to 95°C at 1.0 °C/min for three times.

### Exonuclease digestion assay^43^

For 3’-exonuclease stability, 16.7 μM oligonucleotide (5 nmol) was incubated in a buffer containing 10 mM Tris-HCl (pH 8.0), 2.0 mM MgCl_2_, and 0.2 mU/mL snake venom phosphodiesterase I (SVPD, Aldrich). For 3’-exonuclease stability, 10 μM oligonucleotide was incubated at 37°C in 50 mM NaOAc (pH 6.0) buffer containing 0.25 U/ml bovine spleen phosphodiesterase II (BSP, Aldrich). Right after mixing with enzyme, an aliquot (200 pmol) was collected as an approximate sample of T = ∼0 min. For assay with SVPD, aliquots (200 pmol) of the reaction mixtures were collected at each time point (0.083, 0.15, 0.5, 1, 2, 3, 4, 5, 6, 7, 8, 9, 10, 11, 12, 16, 24, 48 h) and quenched by adding a buffer containing 10 mM EDTA (3x volume of each aliquot) and quickly frozen by liquid nitrogen. For assay with BSP, aliquots (200 pmol) of the reaction mixtures were collected at each time point (0.15, 0.5, 1, 2, 3, 4, 6, 9, 12, 24, 48 h) and quenched by adding a buffer containing EDTA (final EDTA concentration: 2 mM) and quickly frozen by liquid nitrogen. Frozen samples were stored at - 80°C until HPLC analysis. After completion of the incubation, collected samples were heated at 95°C for 15 min, cooled down in ice, spun, and then analyzed by Anion exchange HPLC using Agilent PL-SAX (1000A 8μm, 150×4.6 mm, buffer A: 25 mM sodium phosphate buffer (pH 7.0) at a flow rate 1 mL/min with a gradient mobile phase from 5% buffer B (1 M NaClO_4_ in 25 mM sodium phosphate, pH 7.0) to 40% buffer A (25 mM sodium phosphate, pH 7.0) in 20 min. Oligonucleotide length at each time point was quantified by measuring the area under the curve of the full-length peak and plotted to the graph.

### In-vitro exNA modified siRNA screening

Passive uptake of siRNAs and quantitative analysis of residual nuclear and cytosolic Htt mRNA was performed as previously described^32^. For less efficacious siRNAs, lipid-mediated uptake was performed according to a previously reported protocol^32^.

### Injection of conjugated siRNAs into mice

Animal experiments were performed in accordance with animal care ethics approval and guidelines of the University of Massachusetts Chan Medical School Institutional Animal Care and Use Committee (IACUC, protocol no. A-2411). 6-8-week-old FVB/N female mice (*22–25g) obtained from Charles River (The Jackson Laboratory, Bar Harbor, ME) were administered siRNA by intrascapular subcutaneous injection with 150 μL PBS (PBS controls) or with the indicated amount of siRNA (DCA conjugated) suspended in 150 μL PBS.

### In Vivo mRNA Silencing Experiments

Tissues were collected and stored in RNAlater (Sigma) at 4°C overnight. mRNA was quantified using the QuantiGene 2.0 Assay (Affymetrix). Tissue punches were lysed in 300 μL Homogenizing Buffer (Affymetrix) containing 0.2 mg/mL proteinase K (Invitrogen). Diluted lysates and probe sets (mouse *Htt* or mouse *Hprt*) were added to the bDNA capture plate and the signal was amplified and detected as described previously^60^.

### Peptide Nucleic Acid (PNA) Hybridization Assay for Tissue siRNA Quantification

Quantification of antisense strands in tissues was performed using a PNA hybridization assay as described^61^. Briefly, tissues were lysed in MasterPure tissue lysis solution (EpiCentre) containing 0.2 mg/mL proteinase K (Invitrogen). Sodium dodecyl sulfate (SDS) was precipitated from lysates by adding 3 M potassium chloride and pelleted centrifugation at 4,000 × g for 15 min. siRNA in the supernatant were hybridized to a Cy3-labeled PNA probe fully complementary to the antisense strand (PNABio). Samples were analyzed by HPLC (Agilent) over a DNAPac PA100 anion-exchange column (ThermoFisher). Cy3 fluorescence was monitored, and peaks were integrated. The final concentrations were ascertained using calibration curves.

### Serial blood microsampling

Experiments were conducted as described in the previous paper (Godinho. BMDG. Et al.). Briefly, before the initiation of the experiment, animals were anesthetized, and hair was removed from both hind legs using Nair hair removal cream. siRNA was administered subcutaneously (10 mg/kg) to 6-8-week old FVB/N female mice obtained from Charles River (n = 4/group). Microsamples of blood (10-20 μL) were serially collected from the saphenous vein at time points of 5 min, 15 min, 30 min, 1 h, 3 h, 6 h, 9 h, and 24 h (in a similar manner as described in Ref. J. Pharm. Sci. 2009, 98, 1877–1884). Using a 30G1/2 sterile needle or a lancet, the vein was punctured, and blood droplet was collected using Microvette CB300 K2R tubes (Cat. No. 16.444.100; Sarstedt). Slight pressure is applied on the punctured site to stop the bleeding, and the animal was returned to its cage. Subsequently, collected blood was centrifuged at 10,000 G at 4°C for 15 min to isolate plasma (J Control Release, 2014, 196:106–112. J. Pharm. Sci. 2009, 98, 1877–1884). Plasma was then transferred and stored in Eppendorf tubes at -80°C for subsequent analysis of plasma siRNA concentration. After collection of the last blood sample at 24 h, animals were stored in cage and euthanized at 2 weeks post-injection. Tissues, including liver, kidney, heart, quadriceps muscle, and fat were harvested and placed in RNAlater (Cat. No. R0901; Sigma) overnight. Two 2-mm punches for each tissue were taken, weighed, and lysed as described in the quantitative analysis of target mRNA in vivo. Quantification of oligonucleotides in plasma sample and tissues were conducted by PNA-hybridization assay as described in method. Pharmacokinetic data was fit using a one compartment absorption model using the following equation:

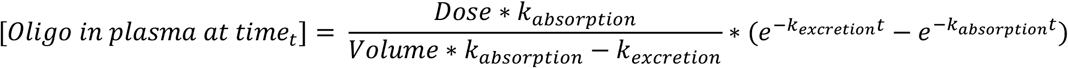

Constants for Dose and Volume were set as:

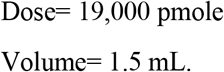

Initial guesses for k_absorption_ and k_excretion_ were both 0.1 and the nonlinear regression function of Graphpad Prism v9.5.1 was used to fit the values for k_absorption_ and k_excretion_.

### Stereotaxic injections in rodents

WT FVB/NJ female mice were purchased from The Jackson Laboratories (strain#001800). All animals were maintained and used according to the Institutional Animal Care and Use Committee guidelines of University of Massachusetts Medical School (docket #20210018). Briefly, mice were housed with a maximum of 5 per cage in a pathogen-free facility under standard conditions with access to food, water, and enrichment *ad libitum*. Mice were injected bilaterally directly into the intracerebroventricular (ICV) system at 8 weeks of age. The mice were anesthetized intraperitoneally using 284 mg/kg tribromoethanol (Avertin) and placed into a small animal stereotaxic apparatus (ASI Instruments and WPI Instruments). Microinjector pumps were fastened to the stereotaxic and set to deliver 5.0μL of the siRNA at a rate of 500 nL/min. Surgical position was set using the bregma as the zero position for guidance, then measured +/- 0.8 mm medial-lateral and -0.2 mm anterior-posterior (AP), and finally lowered -2.5mm dorsal-ventral (DV) into the ventricle. The injection started one minute after the needle was inserted, and the needle was removed one minute after the injection was completed. Once injection was complete, the mice were placed on a heating pad until fully awake and returned to their primary housing facility. Each animal received 0.1cc/10g KETOFEN (ketoprofen) subcutaneously once fully awake. All mice were housed individually post-op to allow healing and avoid losing stitches.

### Rodent tissue collection

For efficacy studies, mice were sacrificed one month after ICV siRNA injections. The mice were deeply anesthetized with tribromoethanol and perfused intracardially with 20mL cold 1X PBS buffer. The brains were immediately sliced into 1.0mm sections using a brain matrix (Kent Scientific Corporation, RBMA-200C). Brain sections floating in cold 1X PBS were punched using 1.5mm sterile biopsy punches. The brain punches were submerged in RNAlater or flash frozen on dry ice and stored at -80°C for mRNA and protein analyses.

### Protein quantification

Protein analysis by capillary immunoassay (Wes system, ProteinSimple) was performed as previously described^62^. Briefly, frozen tissue punches were homogenized in 75 ul 10 mM HEPES pH7.2, 250 mM sucrose, 1 mM EDTA + protease inhibitors (Roche) + 1mM NaF + 1mM Na_3_VO_4_ and sonicated for 10 seconds.

Equal amounts of protein (0.6 ug) were analyzed on the Wes instrument using antibodies to Htt (Ab1, aa1-17, DiFiglia et al., Neuron, 1995) and Vinculin (Sigma V9131) at 1:50 and 1:5000 dilutions respectively. Rabbit and mouse secondaries (ProteinSimple DM-001 and DM-002) were mixed at a 1:1 ratio and the manufacturer’s high molecular weight assay was run. The peak area for Htt normalized to Vinculin was determined using Compass for Simple Western software (ProteinSimple).

### Statistical methods for CNS mRNA and protein silencing data

To assess the average effect of exNA modification, the data was fit to a Gamma family generalized mixed effects linear model with a log link, using the R package glmmTMB^63^. In this model brain region and exNA modification status were used as fixed effects and mouse as a random effect

