## Supporting information for "Extended Nucleic Acid (exNA): A Novel, Biologically Compatible Backbone that Significantly Enhances Oligonucleotide Efficacy *in vivo*"

*[1]RNA Therapeutics Institute, University of Massachusetts Medical School, 368 Plantation Street, Worcester, Massachusetts 01605, United States; [2]Program in Molecular Medicine, University of Massachusetts Medical School; [3]Department of Medicine, University of Massachusetts Medical School; [4] Department of Neurology, Harvard Medical School and Mass General Institute for Neurodegenerative Disease, Charlestown, Massachusetts, United State.*

### Contents

General remarks ---------- S2
**Scheme S1** ---------- S3
Synthesis of compound **5a** ---------- S3
Synthesis of compound **6a** ---------- S3-4
Synthesis of compound **8a** ---------- S4
Synthesis of compound **9a** ---------- S4-5
**Scheme S2** ---------- S5
Synthesis of compound **5b** ---------- S5-6
Synthesis of compound **7b** ---------- S7
Synthesis of compound **8b** ---------- S7-8
Synthesis of compound **9b** ---------- S8
**Scheme S3** ---------- S8
Synthesis of compound **6** ---------- S8-9
Synthesis of compound **7** ---------- S9
Synthesis of compound **8** ---------- S9
Synthesis of compound **9** ---------- S9-10
**Table S1** ---------- S10
**Table S2** ---------- S10
**Table S3** ---------- S11
**Table S4** ---------- S12

**Table S5** ---------------------------------------------------------------------------------------------------------------- S12
**Table S6** ------------------------------------------------------------------------------------------------------------ S12-13
**Table S7** ---------------------------------------------------------------------------------------------------------------- S13
**Table S8** ------------------------------------------------------------------------------------------------------------ S13-14
**Table S9** ---------------------------------------------------------------------------------------------------------------- S14
**Table S10** -------------------------------------------------------------------------------------------------------------- S15
**Table S11** -------------------------------------------------------------------------------------------------------------- S16
**Figure S1** -------------------------------------------------------------------------------------------------------------- S16
**Figure S2** -------------------------------------------------------------------------------------------------------------- S17
**Figure S3** -------------------------------------------------------------------------------------------------------------- S18
**Figure S4** -------------------------------------------------------------------------------------------------------------- S18

### ■ General remarks

The NMR spectra were recorded using Brucker 500 MHz spectrometer. $^{1}$H, $^{13}$C, $^{19}$Fand $^{31}$P NMR spectra were recorded at 500 MHz ($^{1}$H NMR, 500 MHz; $^{13}$C-NMR, 125 MHz; $^{31}$P-NMR, 202 MHz). The chemical shifts were measured from tetramethylsilane (0 ppm), $CDCl_3$ (7.26 ppm), DMSO-$d_6$ (2.49 ppm) and $CD_3CN$-$d_3$ (1.93 ppm) for $^{1}$H-NMR spectra, $CDCl_3$ (77.0 ppm), DMSO-$d_6$ (39.7 ppm) and $CD_3CN$-$d_3$ (1.30 ppm) for $^{13}$C-NMR spectra, and 85% $H_3PO_4$ for $^{31}$P-NMR spectra as external standards. High-resolution electrospray ionization mass spectrometry (HR-ESI-MS) analysis for the monomers and dimer nucleos(t)ides were performed on a Thermo Scientific Orbitrap Velos Pro mass spectrometer in the positive ion mode. The mass analysis of oligonucleotides was conducted by LC-MS on an Agilent 6530 accurate-mass Q-TOF LC/MS (Agilent technologies, Santa Clara, CA). Thin-layer chromatography (TLC) analysis was conducted using silica gel-coated aluminium-backed TLC plates (0.20 mm thickness) containing F-254 UV indicator (Silicycle Inc., Canada). Column chromatography was performed with silica column (Flash Column Silica-CS-Agela; 12-330 g; 40-60 μm), using the CombiFlash Rf200 (Teledyne Isco, Inc.) Companion Chromatograph. The synthesis of modified oligonucleotides was performed using MerMaid-12 DNA/RNA synthesizer (Bio automation, USA). Analytical anion-exchange HPLC and reverse phase HPLC were performed on Agilent 1260 Infinity Analytical SFC System combined with an Agilent 1100 series quaternary pump with a degasser. Purified oligonucleotides were desalted by Sephadex G-25 (GE Healthcare).

**Scheme S1.** 2′-OMe-Ur exNA phosphoramidite

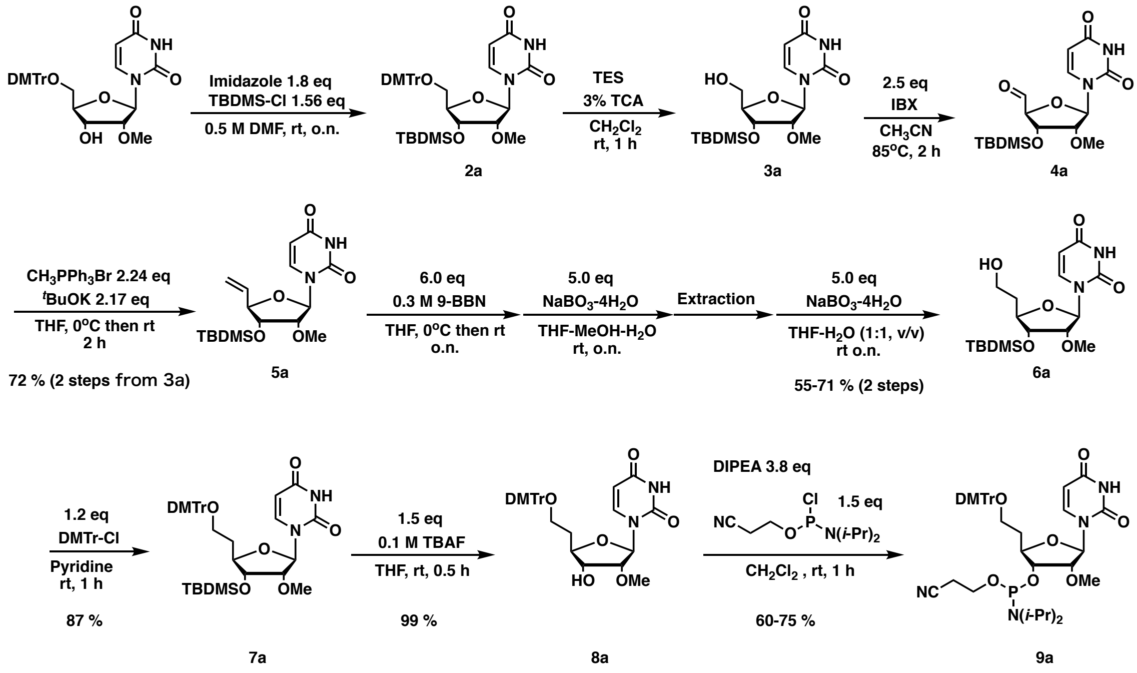


*Synthesis of compound **5a**.* Anhydrous solution of compound **3a** (2.94 g, 7.89 mmol) in $CH_3CN$ (80 mL) was added IBX (5.53 g, 19.7 mmol) and stirred for 2 h at 85 °C. After cooling the mixture in an ice bath, the precipitate in the solution was filtered off through celite. The collected eluent was evaporated, and co-evaporated with anhydrous $CH_3CN$ three times under an argon atmosphere, and obtained compound **4a** as a white foam was used without further purification. In a separate flask, anhydrous THF (80 mL) solution containing methyltriphenylphosphonium bromide (8.47 g, 23.7 mmol) was added *tert*-BuOK (2.57 g, 22.9 mmol) at 0 °C and stirred for 30 min at 0 °C. To this solution, anhydrous THF solution (80 mL) of compound **4a** was added dropwise (10 min) at 0 °C and stirred for 7 h at rt. After evaporating excess THF, the obtained mixture was dissolved in excess ethyl acetate, washed by aq. sat. $NH_4Cl$, dried over $MgSO_4$, filtered, and evaporated. The obtained material was dissolved into a minimum amount of $CH_2Cl_2$ and added dropwise to excess diethyl ether solution under vigorous stirring under the icing. Precipitate in solution was filtered off through celite and eluents was evaporated. The obtained crude material was purified by silica gel column chromatography (hexane/ethyl acetate, 9:1 to 1:2) yielding compound **5a** as a white foam (2.19 g, 75 % in 2 steps). $^1H$ NMR (500 MHz, $CDCl_3$) $\delta$ 9.55 (br-s, 1H), 7.38 (d, 1H, J = 8.2 Hz), 5.89 (ddd, 1H, $J$ = 17.1, 10.6, 6.6 Hz), 5.82 (d, 1H, $J$ = 2.0 Hz), 5.77 (dd, 1H, $J$ = 8.1, 1.5 Hz), 5.44 (dt, 1H, $J$ = 17.2, 1.2 Hz), 5.34 (dt, 1H, $J$ = 10.5, 1.1 Hz), 4.43-4.40 (m, 1H), 3.90 (dd, 1H, $J$ = 7.7, 5.1 Hz), 3.71 (dd, 1H, $J$ = 5.0, 2.0 Hz), 3.55 (s, 3H), 0.89 (s, 9H), 0.09 (s, 3H), 0.07 (s, 3H); $^{13}C$ NMR (125 MHz, $CDCl_3$) $\delta$ 163.4, 150.0, 139.7, 134.4, 119.2, 102.4, 89.7, 84.0, 83.5, 74.5, 58.7, 25.7, 18.2, -4.6, -4.7; HRMS (ESI) calcd. for $C_{17}H_{29}N_2O_5Si^+$ $[M + H]^+$ m/z 369.1840, found m/z 369.1838.

*Synthesis of compound **6a**.* Anhydrous solution of compound **5a** (7.29 g, 19.8 mmol) in THF (158.3 mL) was added 0.5 M 9-BBN/THF solution (237.4 mL, 118.7 mmol) dropwise for 10 min at 0°C. After stirring the mixture at rt 6 h, the solution was iced and added methanol (65.4 mL) and stirred until bubbling cease down. Then under vigorous stirring, $H_2O$ (98.4 mL) was added dropwise for 10 min to avoid precipitation of an intermediate compound. At 0°C, $NaBO_3$-$4H_2O$ (15.7 g, 102.0 mmol) was added in one portion and stirred at rt o.n. After evaporation of excess THF, obtained crude mixture was dissolved into excess ethyl acetate, and washed repeatedly by sat. aq. $NH_4Cl$ solution. After evaporating organic layer, obtained material was dissolved in THF (450 mL) and $H_2O$ (450 mL). To this solution, $NaBO_3$-$4H_2O$ (15.7 g, 102.0 mmol) was added in one portion at rt, then stirred o.n. at rt. After evaporating of excess THF, the mixture was added ethyl acetate, then extracted. Obtained organic layer was repeatedly washed by aq. sat. $NH_4Cl$, dried over $MgSO_4$, filtered, and evaporated. Obtained crude material was purified by silica gel column chromatography (hexane/ethyl acetate, 7:3 to 0:10) yielding compound **6a** as a white foam (4.73 g, 62 % in 2 steps). $^1$H NMR (500 MHz, $CDCl_3$) $\delta$ 9.21(br-s, 1H), 7.35 (d, 1H, J = 8.1 Hz), 5.78-5.76 (m, 2H), 4.14-4.10 (m, 1H), 3.92-3.79 (m, 4H), 3.75 (dd, 1H, J = 5.2, 2.3 Hz), 2.06-2.00 (m, 1H), 1.90-1.82 (m, 1H), 0.91 (s, 9H), 0.11 (s, 3H), 0.10 (s, 3H); $^{13}$C NMR (125 MHz, $CDCl_3$) $\delta$ 163.1, 149.9, 140.0, 102.6, 90.1, 83.0, 82.0, 74.5, 60.3, 58.4, 35.5, 25.7, 18.1, -4.6, -4.9; HRMS (ESI) calcd. for $C_{17}H_{31}N_2O_5Si^+$ $[M + H]^+$ m/z 387.1946, found m/z 387.1944.

*Synthesis of compound **8a**.* Anhydrous solution of compound **6a** (9.46 g, 24.5 mmol) in pyridine (240 mL) was added DMTrCl (9.95 g, 29.4 mmol) and stirred at rt for 2h. After quenching the reaction mixture by MeOH (20 mL), excess pyridine was evaporated, then obtained material was dissolved into excess ethyl acetate. The organic solution was washed by aq. sat. $NaHCO_3$, dried over $MgSO_4$, filtered, evaporated, then co-evaporated with toluene to remove pyridine residues. This crude mixture containing compound **7a** was dissolved into THF (330 mL), added 1.0 M TBAF-THF solution (36.7 mL, 36.7 mmol), then stirred for 1 h at rt. After evaporation excess THF and co-evaporation with $CH_2Cl_2$, the crude material was purified by silica gel column chromatography yielding compound **8a** (13.15 g, 93 % in 2 steps). $^1$H NMR (500 MHz, $CDCl_3$) $\delta$ 8.91 (br-s, 1H), 7.43-7.21 (m, 2H), 7.32-7.14 (m, 8H), 6.83-6.82 (m, 1H), 5.80 (d, 1H, $J$ = 1.8 Hz), 5.69 (d, 1H, $J$ = 8.2 Hz), 4.02-3.98 (m, 1H), 3.85 (dd, 1H, $J$ = 6.7, 6.7 Hz), 3.79 (s, 6H), 3.72 (dd, 1H, J = 5.5, 1.9 Hz), 3.34-3.25 (m, 2H), 2.91 (br-s, 1H), 2.11-2.04 (m, 1H), 1.95-1.89 (m, 1H); $^{13}$C NMR (125 MHz, $CDCl_3$) $\delta$ 163.2, 163.1, 158.4, 149.9, 114.8, 139.1, 136.1, 136.0, 129.92, 129.90, 128.0, 127.8, 126.8, 113.1, 102.5, 88.1, 86.6, 83.5, 81.3, 73.2, 60.1, 58.8, 55.2, 53.4, 33.4; HRMS (ESI) calcd. for $C_{32}H_{34}N_2O_8Na$ $[M + Na]^+$ m/z 597.2203, found m/z 597.2153.

*Synthesis of compound **9a***. Compound **8a** (9.57 g, 16.65 mmol) was rendered anhydrous by repeated co-evaporation with anhydrous $CH_3CN$ and then dissolved into anhydrous $CH_2Cl_2$ (150 mL). To this solution *N*,*N*-diisopropylethylamine (7.6 mL, 62.4 mmol) and 2-cyanoethyl *N*,*N*-diisopropylchlorophosphoramidite (4.85 mL, 25.0 mmol) were added at 0 °C. After stirring for 4 h at rt, the reaction mixture was added $CH_2Cl_2$ (200 mL) then aq. sat. $NaHCO_3$ (350 mL). The organic layer was repeatedly washed by aq. sat. $NaHCO_3$, dried over $MgSO_4$, filtered, then evaporated. The obtained crude material was purified by silica gel column chromatography (1%TEA-hexanes-ethyl acetate, from 80:20 to 30:70) yielding compound **9a** with an impurity of phosphorylating reagent residues. To remove the impurity, obtained material was dissolved in $Et_2O$-ethylacetate (1:1, v/v, 400 mL), then repeatedly washed by aq.

sat. $NaHCO_3$ yielding compound **9a** as a white solid (11.12 g, 86 %); $^{31}P$ NMR (202 MHz, $CDCl_3$) $\delta$ 150.0, 149.9; HRMS (ESI) calcd. for $C_{41}H_{52}N_4O_9P$ $[M + H]^+$ m/z 775.3486, found m/z 775.3414.

**Scheme S2.** Synthesis of 2′-F-Ur exNA phosphoramidite

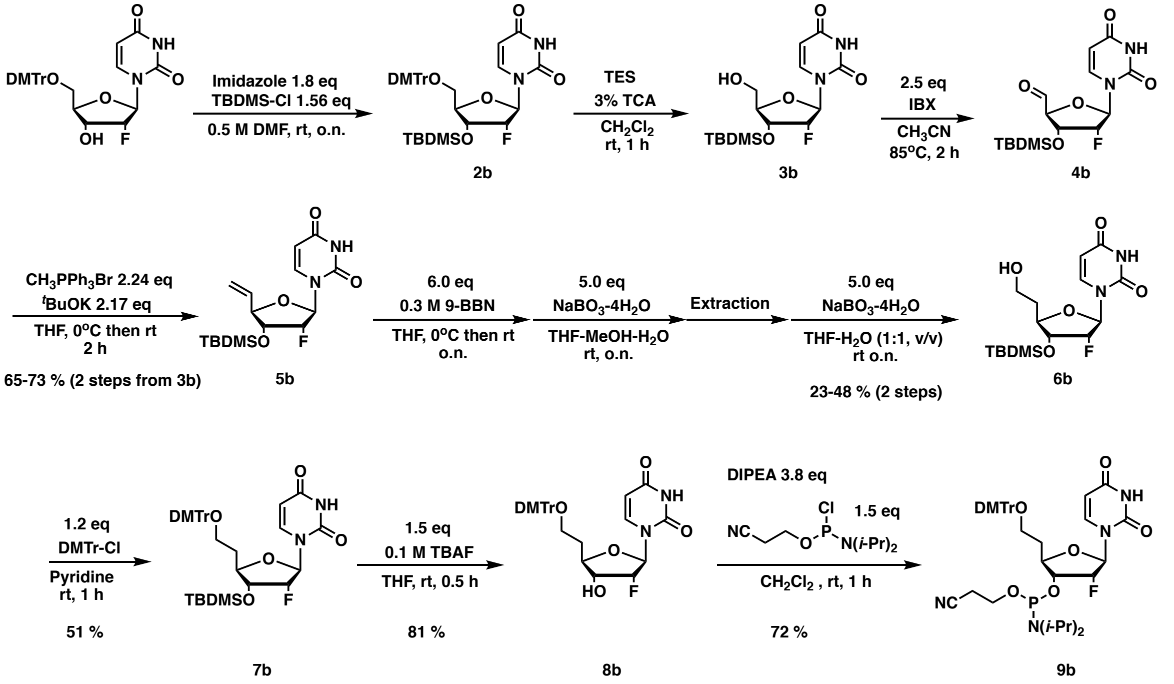


*Synthesis of compound* ***5b*****.** Anhydrous solution of compound **3b** (10.8 g, 30.0 mmol) in $CH_3CN$ (300 mL) was added IBX (21.0 g, 75.0 mmol) and stirred for 2 h at 85 °C. After cooling the mixture in an ice bath, the precipitate in the solution was filtered off through celite. Collected eluent was evaporated, co-evaporated with anhydrous $CH_3CN$ three times under argon atmosphere, and obtained compound **4b** as a white foam was used without further purification. In a separate flask, anhydrous THF (250 mL) solution containing *tert*-BuOK (7.30 g, 65.1 mmol) was added methyltriphenylphosphonium bromide (24.0 g, 68.1 mmol) was added in one portion at 0 °C and stirred for 1 h at 0 °C. To this solution, anhydrous THF solution (150 mL) of compound **4b** was added dropwise (10 min) at 0 °C and stirred o.n. at rt. After evaporating excess THF, the obtained mixture was dissolved in excess ethyl acetate, washed by aq. sat. $NH_4Cl$, dried over $MgSO_4$, filtered, and evaporated. The obtained material was dissolved into minimum amount of $CH_2Cl_2$ and added dropwise to excess diethyl ether solution under vigorous stirring under the icing. Precipitate in the solution was filtered off through celite and eluents were evaporated. Obtained crude material was purified by silica gel column chromatography (hexane/ethyl acetate, 8:2 to 6:4) yielding compound **5b** as a white foam (7.12 g, 67 % in 2 steps). $^1H$ NMR (500 MHz, $CDCl_3$) $\delta$ 11.4 (br-s, 1H), 7.65 (d, 1H, J = 8.1 Hz), 5.93 (ddd, 1H, $J$ = 17.5, 10.4, 7.5 Hz), 5.82 (dd, 1H, $J_{HF}$ = 22.2 Hz, $J_{HH}$= 1.3 Hz), 5.65 (d, 1H, $J$ = 8.1 Hz), 5.42-5.38 (m, 1H), 5.33-5.31 (m, 1H), 5.15 (ddd, 1H, $J_{HF}$ = 53.4 Hz, $J_{HH}$= 4.6, 1.2 Hz), 4.27 (ddd, 1H, $J_{HF}$ = 20.6 Hz, $J_{HH}$ = 8.4, 4.9 Hz), 4.18 (dd, 1H, J = 7.7, 7.7 Hz), 0.88 (s, 9H), 0.08 (s, 3H), 0.07 (s, 3H); $^{13}C$ NMR (125 MHz, $CDCl_3$) $\delta$ 170.8, 183.7, 150.7, 142.5, 135.3, 120.2, 102.4, 92.9 (d, $J_{CF}$ = 186.2 Hz), 90.2 (d, $J_{CF}$ = 36.4 Hz), 83.4, 73.8 (d, $J_{CF}$ = 15.5 Hz), 60.2, 26.0, 21.2, 18.2, 14.6, -4.4, -4.5;

$^{19}F$ NMR (470 MHz, DMSO-*d6*) $\delta$ -198.3 (ddd, $J$ = 53.8, 20.8, 20.8 Hz). HRMS (ESI) calcd. for $C1_6H_{26}FN_2O_4Si$ [M + H]$^+$ m/z 357.1640, found m/z 357.1649.

*Synthesis of compound* **7b.** Anhydrous solution of compound **5b** (10.15 g, 28.5 mmol) in THF (228 mL) was added 0.5 M 9-BBN/THF solution (342 mL, 171 mmol) dropwise for 20 min at 0ºC. After stirring the mixture at rt 4 h, the solution was iced and added methanol (131 mL) and stirred until bubbling cease down. Then under vigorous stirring, $H_2O$ (197 mL) was added dropwise for 15 min to avoid precipitation of an intermediate compound. At 0ºC, $NaBO_3$-$4H_2O$ (21.9 g, 142.5 mmol) was added in one portion and stirred at rt o.n. After evaporation of excess THF, obtained crude mixture was dissolved into excess ethyl acetate, and washed repeatedly by sat. aq. $NH_4Cl$ solution. After evaporating organic layer, obtained material was dissolved in THF (450 mL) and $H_2O$ (450 mL). To this solution, $NaBO_3$-$4H_2O$ (21.9 g, 142.5 mmol) was added in one portion at rt, then stirred o.n. at rt. After evaporating of excess THF, the mixture was added ethyl acetate, then extracted. Obtained organic layer was repeatedly washed by aq. sat. $NH_4Cl$, dried over $MgSO_4$, filtered, and evaporated. Obtained crude material was purified by silica gel column chromatography ($CH_2Cl_2$/methanol, 100:0 to 93:7) yielding compound **6b** as a syrup (2.44 g with reagent impurity); HRMS (ESI) calcd. for $C_{16}H_{28}FN_2O_5Si^+$ [M + H]$^+$ m/z 375.1746, found m/z 375.1746. This compound **6b** with reagent impurity was rendered anhydrous by repeated co-evaporation with anhydrous pyridine under argon atmosphere, then dissolved in anhydrous pyridine (64 mL). To this solution, DMTrCl (2.64 g, 7.79 mmol) was added and stirred at rt for 1 h. After the reaction was quenched by addition of methanol (5 mL), reaction mixture was diluted with ethyl acetate (300 mL) and washed repeated by aq. sat. NaHCO3, dried over MgSO4, filtered, evaporated, then co-evaporated with toluene three times to remove remaining pyridine. Obtained crude material was purified by silica gel chromatography (hexane-ethyl acetate from 2:8 to 4:6) yielding compound **7b** as a white solid (2.25 g, 12% in 2 steps). $^1H$ NMR (500 MHz, $CD_3CN$) $\delta$ 9.16 (br-s, 1H), 7.43-7.42 (m, 2H), 7.31-7.28 (m, 8H), 6.86-6.85 (m, 4H), 5.75 (dd, 1H, $J_{HF}$ = 20.0 Hz, $J_{HH}$ = 1.9 Hz), 5.59 (d, 1H, $J$ = 8.1 Hz), 4.96 (ddd, $J_{HF}$ = 53.3 Hz, $J_{HH}$ = 4.6, 1.8 Hz), 4.06-3.98 (m, 2H), 3.76 (s, 6H), 3.19 (dd, 2H, $J$ = 7.4, 5.6 Hz), 2.09-2.02 (m, 1H), 1.89-1.82 (m, 1H), 0.91 (s, 9H), 0.10 (s, 3H), 0.09 (s, 3H); $^{13}C$ NMR (125 MHz, $CD_3CN$) $\delta$ 163.9, 159.6, 151.1, 146.4, 141.9, 137.31, 137.26, 130.92, 130.89, 128.9, 128.8, 127.8, 114.0, 102.9, 93.7 (d, $J_{CF}$ = 188.0 Hz), 90.6 (d, $J$ = 36.4 Hz), 87.0, 80.7, 74.6 (d, $J$ = 15.4 Hz), 60.9, 55.9, 33.8, 26.1, 18.7, -4.5, -4.8; $^{19}F$ NMR (470 MHz, $CD_3CN$) $\delta$ -201.4 (ddd, $J$ = 53.7, 19.1, 19.1 Hz); HRMS (ESI) calcd. for $C_{37}H_{45}FN_2O_7Na$ [M + Na]$^+$ m/z 699.2872, found m/z 699.2866.

*Synthesis of compound **8b***. Compound 7b (2.24 g, 3.30 mmol) was dissolved into THF (36.0 mL), then added 1.0 M TBAF-THF solution (4.0 mL, 4.0 mmol), then stirred for 30 min at rt. After evaporation excess THF and co-evaporation with $CH_2Cl_2$, the crude material was purified by silica gel column chromatography [$CH_2Cl_2$(1% TEA)-methanol from 100:0 to 95:5] yielding compound **8b** (1.51 g, 81%). $^1H$ NMR (500 MHz, $CDCl_3$) $\delta$ 9.20 (br-s, 1H), 7.45-7.43 (m, 2H), 7.32-7.13 (m, 8H), 6.87-6.85 (m, 4H), 5.77 (dd, 1H, JHF = 20.1 Hz, JHH = 1.5 Hz), 5.60 (d, 1H, J = 8.1 Hz), 4.98 (ddd, JHF= 51.0 Hz, JHH = 4.6, 1.6 Hz), 4.04-3.92 (m, 2H), 3.76 (s, 6H), 3.23-3.17 (m, 2H), 2.20 (br-s, 1H), 2.11-2.06 (m, 1H), 1.91-1.87 (m, 1H); $^{13}C$ NMR (125 MHz, $CD_3CN$) $\delta$ 164.1, 159.7, 151.2, 146.4 141.7, 139.0,

137.33, 137.29, 131,00, 130.97, 129.3, 129.0, 128.9, 127.8, 126.3, 114.1, 103.0, 94.7 (d, J = 184.4 Hz), 90.3 (d, J = 35.4 Hz), 87.2, 80.5, 73.9 (d, J = 16.4 Hz), 61.0, 56.0, 33.9; $^{19}$F NMR (470 MHz, $CD_3CN$) $\delta$ -201.8 (ddd, J = 53.7, 20.8, 20.8 Hz). HRMS (ESI) calcd. for $C_{31}H_{31}FN_2O_7Na$ $[M + Na]^+$ m/z 585.2013, found m/z 585.1999.

*Synthesis of compound **9b***. Compound **8b** (1.5 g, 2.67 mmol) was rendered anhydrous by repeated co-evaporation with anhydrous $CH_3CN$ and then dissolved into anhydrous $CH_2Cl_2$ (30 mL). To this solution *N*,*N*-diisopropylethylamine (1.76 mL, 10.1 mmol) and 2-cyanoethyl *N*,*N*-diisopropylchlorophosphoramidite (0.90 mL, 4.01 mmol) were added at 0 °C. After stirring for 2 h at rt, the reaction mixture was added $CH_2Cl_2$ (70 mL) then aq. sat. $NaHCO_3$ (100 mL). Organic layer was repeatedly washed by aq. sat. $NaHCO_3$, dried over $MgSO_4$, filtered, then evaporated. Obtained crude material was purified by silica gel column chromatography (1%TEA-hexanes-ethyl acetate, from 80:20 to 20:80) yielding compound **9a** with an impurity of phosphitylating reagent residues. To remove the impurity, obtained material was dissolved in $Et_2O$ (100 mL), then repeatedly washed by aq. sat. $NaHCO_3$ yielding compound **9b** as a white solid (1.47 g, 64 %); $^{31}$P NMR (202 MHz, $CDCl_3$) $\delta$ 150.4 (d, *J* = 9.0 Hz), 149.9 (d, *J* = 10.0 Hz); $^{19}$F NMR (470 MHz, $CD_3CN$) $\delta$ -198.61, -198.63, -198.66, -198.68, -198.70, -198.73, -198.75, -198.77, -198.79, -198.82, -198.84, -199.04, -199.06, -199.08, -199.10, -199.12, -199.14, -199.15, -199.17, -199.20, -199.21, -199.24, -199.26. HRMS (ESI) calcd. for $C_{40}H_{49}FN_4O_8P$ $[M + H]^+$ m/z 763.3267, found m/z 763.3255.

**Scheme S3**. Synthesis of 2′-riboUr-exNA phosphoramidite

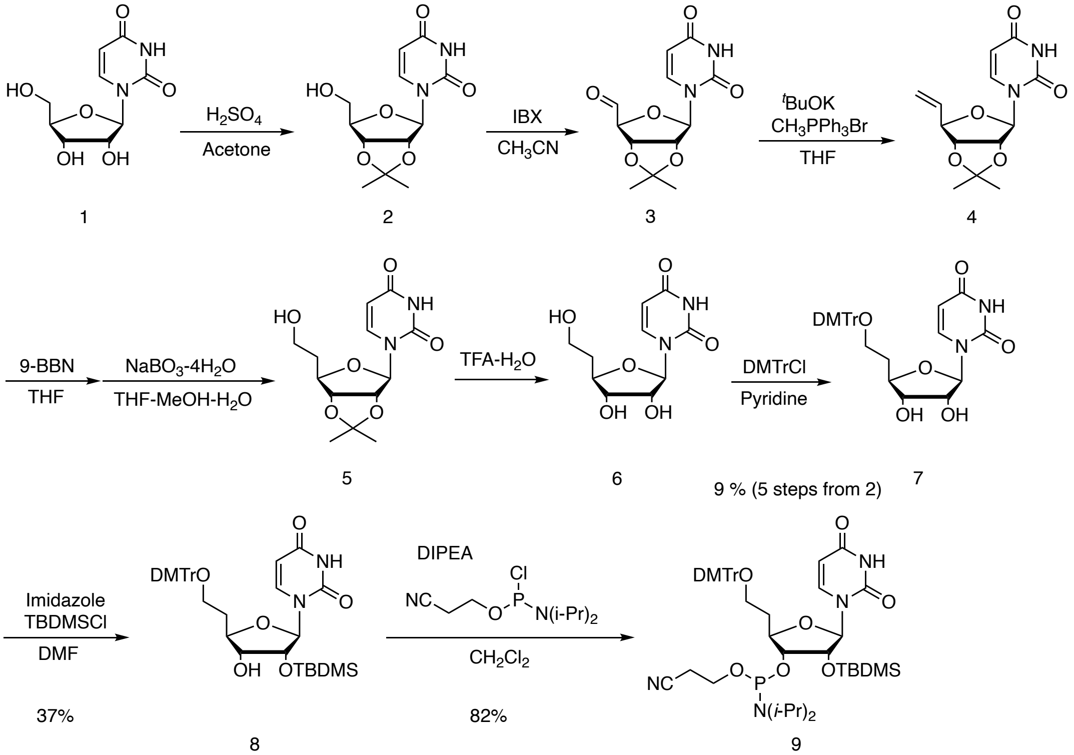


*Synthesis of compound **6***. Anhydrous solution of compound **2** (15.4 g, 54.1 mmol) in $CH_3CN$ (520 mL) was added IBX (30.3 g, 108.2 mmol) and stirred for 2 h at 85 °C. After cooling the mixture in an ice bath, the precipitate in the solution was filtered off through celite. Collected eluent was evaporated, co-evaporated with anhydrous $CH_3CN$ three times under argon atmosphere, and obtained compound **3** as a white foam was used without further purification. In a separate flask, anhydrous THF (500 mL) solution containing *tert*-BuOK (13.2 g, 117.4 mmol) was added methyltriphenylphosphonium bromide (43.3 g, 121.2 mmol) was added in one portion at 0 °C and stirred for 1 h at 0 °C. To this solution, anhydrous THF solution (150 mL) of compound **3** was added dropwise (10 min) at 0 °C and stirred for 4 h. at rt. After evaporating excess THF, the obtained mixture was dissolved in excess ethyl acetate, washed by aq. sat. $NH_4Cl$, dried over $MgSO_4$, filtered, and evaporated. Obtained material was dissolved into minimum amount of $CH_2Cl_2$ and added dropwise to excess diethyl ether solution under vigorously stirring at 0 °C. Precipitate in solution was filtered off through celite and eluents was evaporated. Obtained crude material was purified by silica gel column chromatography (hexane/ethyl acetate, 8:2 to 3:7) yielding compound **4** with impurity of triphenylphosphineoxide. 3/4 of this crude material was rendered anhydrous by repeated co-evaporation with anhydrous $CH_3CN$, and then dissolved in anhydrous THF (200 mL). To this solution, 0.5 M 9-BBN/THF (300 mL, 150.0 mmol) was added dropwisely for 10 min, then stirred at rt o.n. After confirming disappearance of starting material by TLC, the solution was iced, then added methanol (200 mL) dropwisely for 10 min. After bubbling is cease down, $H_2O$ (300 mL) was added dropwise then $NaBO_3$-4H2O (19.2 g, 125.0 mmol) was added in one portion. The solution was stirred o.n. at rt. After evaporation of excess THF, obtained crude mixture was dissolved into excess ethyl acetate, and washed repeatedly by sat. aq. $NH_4Cl$ solution. After evaporating organic layer, obtained material was dissolved in THF (400 mL) and $H_2O$ (400 mL).

To this solution, $NaBO_3$-$4H_2O$ (19.2 g, 125.0 mmol) was added in one portion at rt, then stirred o.n. at rt. After evaporating of excess THF, the mixture was added ethyl acetate, then extracted. Obtained organic layer was repeatedly washed by aq. sat. $NH_4Cl$, dried over $MgSO_4$, filtered, and evaporated. Obtained crude material was purified by silica gel column chromatography ($CH_2Cl_2$-methanol, 100: 0 to 93:7) yielding compound **5** with impurity of reagent residues. This obtained material was added TFA solution [TFA (85 mL) and H2O (9.2 mL)] and stirred at 0ºC for 1 h. After evaporation, co-evaporation with toluene four times, crude material was purified by silica gel column chromatography ($CH_2Cl_2$-MeOH from 100:0 to 90:10) yielding compound **6** (760 mg, 12% in 3 steps). $^1$H NMR (500 MHz, DMSO-d6) $\delta$ 11.4 (br-s, 1H), 7.58 (d, 1H, J = 5.0 Hz), 5.71 (d, 1H, J = 5.0 Hz), 5.64 (dd, 1H, J = 8.0, 2.2 Hz), 5.34 (d, 1H, J = 5.2 Hz), 5.09 (d, 1H, J = 4.7 Hz), 4.51 (br-s, 1H), 4.06, (dd, 1H, J = 9.8, 4.9 Hz), 3.80-3.78 (m, 1H), 3.53-3.45 (m, 2H), 1.84-1.70 (m, 2H); $^{13}$C NMR (125 MHz, DMSO-d6) $\delta$ 163.5, 151.1, 141.6, 102.5, 89.0, 80.9, 73.5, 73.2, 58.0, 46.2, 36.8, 9.1; HRMS (ESI) calcd. for $C_{10}H_{14}N_2O_6Na$ $[M + Na]^+$ m/z 281.0744, found m/z 281.0730.

*Synthesis of compound 7.* The compound **6** (760 mg, 2.94 mmol) was added anhydrous pyridine (30 mL) and then added DMTr-Cl (1.3 g, 3.82 mmol). After stirring for 2 h, reaction mixture was first extracted with $CH_2Cl_2$ an aq. sat. $NaHCO_3$, and organic layer was dried over $MgSO_4$, filtered, evaporated, co-evaporated to remove pyridine. The obtained crude material was used for the next reaction without further purification. HRMS (ESI) calcd. for $C_{31}H_{32}N_2O_8Na$ $[M + Na]^+$ m/z 583.2051, found m/z 583.2025.

*Synthesis of compound **8***. An anhydrous solution of compound **7** (2.35 g, 4.19 mmol) in pyridine (21 mL) was added imidazole (576.1 mg, 8.46 mmol) and TBDMSCl (1.10 g, 7.33 mmol), and then stirred for 2h at rt. To this reaction mixture was added $CH_2Cl_2$ (150 mL) was then added aq. sat. $NaHCO_3$ (150 mL). The organic layer was repeatedly washed by aq. sat. $NaHCO_3$, dried over $MgSO_4$, filtered, evaporated, then co-evaporated with toluene to remove pyridine residue. Obtained crude material containing compound 8, 3'-O-TBDMS protected compound, 5'-3'-O-bis-TBDMS protected compound was separated by silica gel column chromatography [$CH_2Cl_2$ (1% TEA)-Acetone from 100:0 to 85:15] yielding pure compound 8 (780 mg, 28%). $^1$H NMR (500 MHz, DMSO-d6) $\delta$ 11.4 (br-s, 1H), 7.53-7.52 (m, 2H), 7.39-7.22 (m, 8H), 6.89-6.88 (m, 4H), 5.72 (d, 1H, J = 5.0 Hz), 5.62 (d, 1H, J = 8.1, 2.0 Hz), 5.00 (d, 1H, J = 6.0 Hz), 4.19 (dd, 1H, J = 5.1, 5.1 Hz), 3.92 (ddd, 1H, J = 8.8, 8.8, 4.5 Hz), 3.78-3.73 (m, 7H), 3.07-3.03 (m, 2H), 2.05-1.83 (m, 2H), 0.83 (s, 9H), 0.05 (s, 3H), 0.01 (s, 3H); $^{13}$C NMR (125 MHz, DMSO-$d_6$) $\delta$ 162.9, 158.0, 150.5, 145.1, 140.7, 135.8, 130.1, 129.4, 128.7, 128.3, 128.1, 127.1, 125.8, 113.6, 102.5, 88.8, 86.0, 81.5, 74.9, 73.4, 60.6, 55.5, 33.8, 26.1, 25.1, 18.4; HRMS (ESI) calcd. for $C_{37}H_{46}N_2O_8Na$ $[M + Na]^+$ m/z 697.2916, found m/z 697.2867.

*Synthesis of compound **9***. Compound **8** (780 g, 1.16 mmol) was rendered anhydrous by repeated co-evaporation with anhydrous $CH_3CN$ and then dissolved into anhydrous $CH_2Cl_2$ (12 mL). To this solution *N*,*N*-diisopropylethylamine

(0.53 mL, 4.34 mmol) and 2-cyanoethyl *N*,*N*-diisopropylchlorophosphoramidite (0.34 mL, 1.73 mmol) were added at 0 °C. After stirring for 4 h at rt, the reaction mixture was added $CH_2Cl_2$ (90 mL) then aq. sat. $NaHCO_3$ (100 mL). Organic layer was repeatedly washed by aq. sat. $NaHCO_3$, dried over $MgSO_4$, filtered, then evaporated. Obtained crude material was purified by silica gel column chromatography (1%TEA-hexanes-ethyl acetate, from 80:20 to 50:50) yielding compound 9 (825.9 mg, 82%). $^{31}P$ NMR (202 MHz, $CDCl_3$) $\delta$ 149.6, 149.1.

**Table S1**. List of exNA–modified siRNA duplexes used for in–vitro screening.

| Duplex# | Guide strand | Sequence (5' -> 3') | Passenger strand | Sequence (5' -> 3') |
|---|---|---|---|---|
| **D1** | ex-AS-1 | P(**ex_mU**)#(fU)#(mA)(fA)(mU)(fC)(mU)(fC)(mU)(fU)(mU)(fA)(mC)#(fU)#(mG)#(fA)#(mU)#(fA)#(mU)#(fA) | SS-1 | (fC)#(mA)#(fG)(mU)(fA)(mA)(fA)(mG)(fA)(mG)(fA)(mU)(fU)#(mA)#(fA)-TegChol |
| **D2** | ex-AS-2 | P(mU)#(**ex_fU**)#(mA)(fA)(mU)(fC)(mU)(fC)(mU)(fU)(mU)(fA)(mC)#(fU)#(mG)#(fA)#(mU)#(fA)#(mU)#(fA) | SS-1 | (fC)#(mA)#(fG)(mU)(fA)(mA)(fA)(mG)(fA)(mG)(fA)(mU)(fU)#(mA)#(fA)-TegChol |
| **D3** | ex-AS-3 | P(mU)#(fU)#(**ex_mU**)(fU)(mU)(fA)(mA)(fA)(mU)(fC)(mC)(fU)(mG)#(fA)#(mG)#(fA)#(mA)#(fG)#(mA)#(fA) | SS-2 | (fC)#(mU)#(fC)(mA)(fG)(mG)(fA)(mU)(fU)(mU)(fA)(mA)(fA)#(mA)#(fA)-TegChol |
| **D4** | ex-AS-4 | P(mU)#(fU)#(mU)(**ex_fU**)(mU)(fA)(mA)(fA)(mU)(fC)(mC)(fU)(mG)#(fA)#(mG)#(fA)#(mA)#(fG)#(mA)#(fA) | SS-2 | (fC)#(mU)#(fC)(mA)(fG)(mG)(fA)(mU)(fU)(mU)(fA)(mA)(fA)#(mA)#(fA)-TegChol |
| **D5** | ex-AS-5 | P(mU)#(fU)#(mA)(fA)(**ex_mU**)(fC)(mU)(fC)(mU)(fU)(mU)(fA)(mC)#(fU)#(mG)#(fA)#(mU)#(fA)#(mU)#(fA) | SS-1 | (fC)#(mA)#(fG)(mU)(fA)(mA)(fA)(mG)(fA)(mG)(fA)(mU)(fU)#(mA)#(fA)-TegChol |
| **D6** | ex-AS-6 | P(mU)#(fC)#(mC)(fA)(mC)(**ex_fU**)(mA)(fU)(mG)(fU)(mU)(fU)(mU)#(fC)#(mA)#(fC)#(mA)#(fU)#(mA)#(fU) | SS-3 | (fU)#(mG)#(fA)(mA)(fA)(mA)(fC)(mA)(fU)(mA)(fG)(mU)(fG)#(mG)#(fA)-TegChol |
| **D7** | ex-AS-7 | P(mU)#(fU)#(mA)(fA)(mU)(fC)(**ex_mU**)(fC)(mU)(fU)(mU)(fA)(mC)#(fU)#(mG)#(fA)#(mU)#(fA)#(mU)#(fA) | SS-1 | (fC)#(mA)#(fG)(mU)(fA)(mA)(fA)(mG)(fA)(mG)(fA)(mU)(fU)#(mA)#(fA)-TegChol |
| **D8** | ex-AS-8 | P(mU)#(fC)#(mC)(fA)(mC)(fU)(mA)(**ex_fU**)(mG)(fU)(mU)(fU)(mU)#(fC)#(mA)#(fC)#(mA)#(fU)#(mA)#(fU) | SS-3 | (fU)#(mG)#(fA)(mA)(fA)(mA)(fC)(mA)(fU)(mA)(fG)(mU)(fG)#(mG)#(fA)-TegChol |
| **D9** | ex-AS-9 | P(mU)#(fU)#(mA)(fA)(mU)(fC)(mU)(fC)(**ex_mU**)(fU)(mU)(fA)(mC)#(fU)#(mG)#(fA)#(mU)#(fA)#(mU)#(fA) | SS-1 | (fC)#(mA)#(fG)(mU)(fA)(mA)(fA)(mG)(fA)(mG)(fA)(mU)(fU)#(mA)#(fA)-TegChol |
| **D10** | ex-AS-10 | P(mU)#(fU)#(mA)(fA)(mU)(fC)(mU)(fC)(mU)(**ex_fU**)(mU)(fA)(mC)#(fU)#(mG)#(fA)#(mU)#(fA)#(mU)#(fA) | SS-1 | (fC)#(mA)#(fG)(mU)(fA)(mA)(fA)(mG)(fA)(mG)(fA)(mU)(fU)#(mA)#(fA)-TegChol |
| **D11** | ex-AS-11 | P(mU)#(fU)#(mA)(fA)(mU)(fC)(mU)(fC)(mU)(fU)(**ex_mU**)(fA)(mC)#(fU)#(mG)#(fA)#(mU)#(fA)#(mU)#(fA) | SS-1 | (fC)#(mA)#(fG)(mU)(fA)(mA)(fA)(mG)(fA)(mG)(fA)(mU)(fU)#(mA)#(fA)-TegChol |
| **D12** | ex-AS-12 | P(mU)#(fC)#(mC)(fA)(mC)(fU)(mA)(fU)(mG)(fU)(mU)(**ex_fU**)(mU)#(fC)#(mA)#(fC)#(mA)#(fU)#(mA)#(fU) | SS-3 | (fU)#(mG)#(fA)(mA)(fA)(mA)(fC)(mA)(fU)(mA)(fG)(mU)(fG)#(mG)#(fA)-TegChol |
| **D13** | ex-AS-13 | P(mU)#(fC)#(mC)(fA)(mC)(fU)(mA)(fU)(mG)(fU)(mU)(fU)(**ex_mU**)#(fC)#(mA)#(fC)#(mA)#(fU)#(mA)#(fU) | SS-3 | (fU)#(mG)#(fA)(mA)(fA)(mA)(fC)(mA)(fU)(mA)(fG)(mU)(fG)#(mG)#(fA)-TegChol |
| **D14** | ex-AS-14 | P(mU)#(fU)#(mA)(fA)(mU)(fC)(mU)(fC)(mU)(fU)(mU)(fA)(mC)#(**ex_fU**)#(mG)#(fA)#(mU)#(fA)#(mU)#(fA) | SS-1 | (fC)#(mA)#(fG)(mU)(fA)(mA)(fA)(mG)(fA)(mG)(fA)(mU)(fU)#(mA)#(fA)-TegChol |
| **D15** | ex-AS-15 | P(mU)#(fG)#(mC)(fC)(mU)(fA)(mA)(fG)(mA)(fG)(mC)(fA)(mC)#(fA)#(**ex_mU**)#(fU)#(mU)#(fA)#(mG)#(fU) | SS-4 | (fA)#(mU)#(fG)(mU)(fG)(mC)(fU)(mC)(fU)(mU)(fA)(mG)(fG)#(mC)#(fA)-TegChol |
| **D16** | ex-AS-16 | P(mU)#(fG)#(mC)(fC)(mU)(fA)(mA)(fG)(mA)(fG)(mC)(fA)(mC)#(fA)#(mU)#(**ex_fU**)#(mU)#(fA)#(mG)#(fU) | SS-4 | (fA)#(mU)#(fG)(mU)(fG)(mC)(fU)(mC)(fU)(mU)(fA)(mG)(fG)#(mC)#(fA)-TegChol |
| **D17** | ex-AS-17 | P(mU)#(fU)#(mA)(fA)(mU)(fC)(mU)(fC)(mU)(fU)(mU)(fA)(mC)#(fU)#(mG)#(fA)#(**ex_mU**)#(fA)#(mU)#(fA) | SS-1 | (fC)#(mA)#(fG)(mU)(fA)(mA)(fA)(mG)(fA)(mG)(fA)(mU)(fU)#(mA)#(fA)-TegChol |
| **D18** | ex-AS-18 | P(mU)#(fC)#(mC)(fA)(mC)(fU)(mA)(fU)(mG)(fU)(mU)(fU)(mU)#(fC)#(mA)#(fC)#(mA)#(**ex_fU**)#(mA)#(fU) | SS-3 | (fU)#(mG)#(fA)(mA)(fA)(mA)(fC)(mA)(fU)(mA)(fG)(mU)(fG)#(mG)#(fA)-TegChol |
| **D19** | ex-AS-19 | P(mU)#(fU)#(mA)(fA)(mU)(fC)(mU)(fC)(mU)(fU)(mU)(fA)(mC)#(fU)#(mG)#(fA)#(mU)#(fA)#(**ex_mU**)#(fA) | SS-1 | (fC)#(mA)#(fG)(mU)(fA)(mA)(fA)(mG)(fA)(mG)(fA)(mU)(fU)#(mA)#(fA)-TegChol |
| **D20** | ex-AS-20 | P(mU)#(fC)#(mC)(fA)(mC)(fU)(mA)(fU)(mG)(fU)(mU)(fU)(mU)#(fC)#(mA)#(fC)#(mA)#(fU)#(mA)#(**ex_fU**) | SS-3 | (fU)#(mG)#(fA)(mA)(fA)(mA)(fC)(mA)(fU)(mA)(fG)(mU)(fG)#(mG)#(fA)-TegChol |
| **D21** | ex-AS-21 | P(mU)#(fU)#(mA)(fA)(mU)(fC)(mU)(fC)(mU)(fU)(mU)(fA)(mC)#(fU)#(mG)#(fA)#(mU)#(fU)#(mU)#(**ex_fU**) | SS-1 | (fC)#(mA)#(fG)(mU)(fA)(mA)(fA)(mG)(fA)(mG)(fA)(mU)(fU)#(mA)#(fA)-TegChol |
| **D22** | ex-AS-22 | P(mU)#(fU)#(mA)(fA)(mU)(fC)(mU)(fC)(mU)(fU)(mU)(fA)(mC)#(fU)#(mG)#(fA)#(mU)#(fU)#(**ex_mU**)#(**ex_fU**) | SS-1 | (fC)#(mA)#(fG)(mU)(fA)(mA)(fA)(mG)(fA)(mG)(fA)(mU)(fU)#(mA)#(fA)-TegChol |
| **D23** | ex-AS-23 | P(mU)#(fU)#(mA)(fA)(mU)(fC)(mU)(fC)(mU)(fU)(mU)(fA)(mC)#(fU)#(mG)#(fA)#(mU)#(**ex_fU**)#(**ex_mU**)#(**ex_fU**) | SS-1 | (fC)#(mA)#(fG)(mU)(fA)(mA)(fA)(mG)(fA)(mG)(fA)(mU)(fU)#(mA)#(fA)-TegChol |
| **D24** | ex-AS-24 | P(mU)#(fU)#(mA)(fA)(mU)(fC)(mU)(fC)(mU)(fU)(mU)(fA)(mC)#(fU)#(mG)#(fA)#(**ex_mU**)#(**ex_fU**)#(**ex_mU**)#(**ex_fU**) | SS-1 | (fC)#(mA)#(fG)(mU)(fA)(mA)(fA)(mG)(fA)(mG)(fA)(mU)(fU)#(mA)#(fA)-TegChol |
| **D25** | AS-1 | P(mU)#(fC)#(mC)(fA)(mC)(fU)(mA)(fU)(mG)(fU)(mU)(fU)(mU)#(fC)#(mA)#(fC)#(mA)#(fU)#(mA)#(fU) | ex-SS-1 | (**ex_fU**)#(mG)#(fA)(mA)(fA)(mA)(fC)(mA)(fU)(mA)(fG)(mU)(fG)#(mG)#(fA)-TegChol |
| **D26** | AS-2 | P(mU)#(fU)#(mU)(fU)(mU)(fA)(mA)(fA)(mU)(fC)(mC)(fU)(mG)#(fA)#(mG)#(fA)#(mA)#(fG)#(mA)#(fA) | ex-SS-2 | (fC)#(**ex_mU**)#(fC)(mA)(fG)(mG)(fA)(mU)(fU)(mU)(fA)(mA)(fA)#(mA)#(fA)-TegChol |
| **D27** | AS-3 | P(mU)#(fC)#(mC)(fG)(mG)(fU)(mC)(fA)(mC)(fA)(mA)(fC)(mA)#(fU)#(mU)#(fG)#(mU)#(fG)#(mG)#(fU) | ex-SS-3 | (fA)#(mA)#(**ex_fU**)(mG)(fU)(mU)(fG)(mU)(fG)(mA)(fC)(mC)(fG)#(mG)#(fA)-TegChol |
| **D28** | AS-4 | P(mU)#(fU)#(mA)(fA)(mU)(fC)(mU)(fC)(mU)(fU)(mU)(fA)(mC)#(fU)#(mG)#(fA)#(mU)#(fA)#(mU)#(fA) | ex-SS-4 | (fC)#(mA)#(fG)(**ex_mU**)(fA)(mA)(fA)(mG)(fA)(mG)(fA)(mU)(fU)#(mA)#(fA)-TegChol |
| **D29** | AS-3 | P(mU)#(fC)#(mC)(fG)(mG)(fU)(mC)(fA)(mC)(fA)(mA)(fC)(mA)#(fU)#(mU)#(fG)#(mU)#(fG)#(mG)#(fU) | ex-SS-5 | (fA)#(mA)#(fU)(mG)(**ex_fU**)(mU)(fG)(mU)(fG)(mA)(fC)(mC)(fG)#(mG)#(fA)-TegChol |
| **D30** | AS-3 | P(mU)#(fC)#(mC)(fG)(mG)(fU)(mC)(fA)(mC)(fA)(mA)(fC)(mA)#(fU)#(mU)#(fG)#(mU)#(fG)#(mG)#(fU) | ex-SS-6 | (fA)#(mA)#(fU)(mG)(fU)(**ex_mU**)(fG)(mU)(fG)(mA)(fC)(mC)(fG)#(mG)#(fA)-TegChol |
| **D31** | AS-5 | P(mU)#(fG)#(mC)(fC)(mU)(fA)(mA)(fG)(mA)(fG)(mC)(fA)(mC)#(fA)#(mU)#(fU)#(mU)#(fA)#(mG)#(fU) | ex-SS-7 | (fA)#(mU)#(fG)(mU)(fG)(mC)(**ex_fU**)(mC)(fU)(mU)(fA)(mG)(fG)#(mC)#(fA)-TegChol |
| **D32** | AS-2 | P(mU)#(fU)#(mU)(fU)(mU)(fA)(mA)(fA)(mU)(fC)(mC)(fU)(mG)#(fA)#(mG)#(fA)#(mA)#(fG)#(mA)#(fA) | ex-SS-8 | (fC)#(mU)#(fC)(mA)(fG)(mG)(fA)(**ex_mU**)(fU)(mU)(fA)(mA)(fA)#(mA)#(fA)-TegChol |
| **D33** | AS-2 | P(mU)#(fU)#(mU)(fU)(mU)(fA)(mA)(fA)(mU)(fC)(mC)(fU)(mG)#(fA)#(mG)#(fA)#(mA)#(fG)#(mA)#(fA) | ex-SS-9 | (fC)#(mU)#(fC)(mA)(fG)(mG)(fA)(mU)(**ex_fU**)(mU)(fA)(mA)(fA)#(mA)#(fA)-TegChol |
| **D34** | AS-2 | P(mU)#(fU)#(mU)(fU)(mU)(fA)(mA)(fA)(mU)(fC)(mC)(fU)(mG)#(fA)#(mG)#(fA)#(mA)#(fG)#(mA)#(fA) | ex-SS-10 | (fC)#(mU)#(fC)(mA)(fG)(mG)(fA)(mU)(fU)(**ex_mU**)(fA)(mA)(fA)#(mA)#(fA)-TegChol |
| **D35** | AS-6 | P(mU)#(fA)#(mU)(fC)(mA)(fG)(mC)(fU)(mU)(fU)(mU)(fC)(mC)#(fA)#(mG)#(fG)#(mG)#(fU)#(mC)#(fG) | ex-SS-11 | (fC)#(mU)#(fG)(mG)(fA)(mA)(fA)(mA)(fG)(mC)(**ex_fU**)(mG)(fA)#(mU)#(fA)-TegChol |
| **D36** | AS-4 | P(mU)#(fU)#(mA)(fA)(mU)(fC)(mU)(fC)(mU)(fU)(mU)(fA)(mC)#(fU)#(mG)#(fA)#(mU)#(fA)#(mU)#(fA) | ex-SS-12 | (fC)#(mA)#(fG)(mU)(fA)(mA)(fA)(mG)(fA)(mG)(fA)(**ex_mU**)(fU)#(mA)#(fA)-TegChol |
| **D37** | AS-4 | P(mU)#(fU)#(mA)(fA)(mU)(fC)(mU)(fC)(mU)(fU)(mU)(fA)(mC)#(fU)#(mG)#(fA)#(mU)#(fA)#(mU)#(fA) | ex-SS-13 | (fC)#(mA)#(fG)(mU)(fA)(mA)(fA)(mG)(fA)(mG)(fA)(mU)(**ex_fU**)#(mA)#(fA)-TegChol |
| **D38** | AS-6 | P(mU)#(fA)#(mU)(fC)(mA)(fG)(mC)(fU)(mU)(fU)(mU)(fC)(mC)#(fA)#(mG)#(fG)#(mG)#(fU)#(mC)#(fG) | ex-SS-14 | (fC)#(mU)#(fG)(mG)(fA)(mA)(fA)(mA)(fG)(mC)(fU)(mG)(fA)#(**ex_mU**)#(fA)-TegChol |
| **D39** | AS-7 | P(mA)#(fU)#(mA)(fA)(mU)(fC)(mU)(fC)(mU)(fU)(mU)(fA)(mC)#(fU)#(mG)#(fA)#(mU)#(fA)#(mU)#(fA) | ex-SS-15 | (fC)#(mA)#(fG)(mU)(fA)(mA)(fA)(mG)(fA)(mG)(fA)(mU)(fU)#(mA)#(**ex_fU**)-TegChol |

**Table S2**. List of control siRNA duplexes used for in-vitro screening.

| | | | | |
|---|---|---|---|---|
| **D40** | **AS-4** | P(mU)#(fU)#(mA)(fA)(mU)(fC)(mU)(fC)(mU)(fU)(mU)(fA)(mC)#(fU)#(mG)#(fA)#(mU)#(fA)#(mU)#(fA) | **SS-1** | (fC)#(mA)#(fG)(mU)(fA)(mA)(fA)(mG)(fA)(mG)(fA)(mU)(fU)#(mA)#(fA)-TegChol |
| **D41** | **AS-8** | P(mU)#(fU)#(mA)(fA)(mU)(fC)(mU)(fC)(mU)(fU)(mU)(fA)(mC)#(fU)#(mG)#(fA)#(mU)#(fU)#(mU)#(fU) | **SS-1** | (fC)#(mA)#(fG)(mU)(fA)(mA)(fA)(mG)(fA)(mG)(fA)(mU)(fU)#(mA)#(fA)-TegChol |
| **D42** | **AS-2** | P(mU)#(fU)#(mU)(fU)(mU)(fA)(mA)(fA)(mU)(fC)(mC)(fU)(mG)#(fA)#(mG)#(fA)#(mA)#(fG)#(mA)#(fA) | **SS-2** | (fC)#(mU)#(fC)(mA)(fG)(mG)(fA)(mU)(fU)(mU)(fA)(mA)(fA)#(mA)#(fA)-TegChol |
| **D43** | **AS-1** | P(mU)#(fC)#(mC)(fA)(mC)(fU)(mA)(fU)(mG)(fU)(mU)(fU)(mU)#(fC)#(mA)#(fC)#(mA)#(fU)#(mA)#(fU) | **SS-3** | (fU)#(mG)#(fA)(mA)(fA)(mA)(fC)(mA)(fU)(mA)(fG)(mU)(fG)#(mG)#(fA)-TegChol |
| **D44** | **AS-5** | P(mU)#(fG)#(mC)(fC)(mU)(fA)(mA)(fG)(mA)(fG)(mC)(fA)(mC)#(fA)#(mU)#(fU)#(mU)#(fA)#(mG)#(fU) | **SS-4** | (fA)#(mU)#(fG)(mU)(fG)(mC)(fU)(mC)(fU)(mU)(fA)(mG)(fG)#(mC)#(fA)-TegChol |
| **D45** | **AS-6** | P(mU)#(fA)#(mU)(fC)(mA)(fG)(mC)(fU)(mU)(fU)(mU)(fC)(mC)#(fA)#(mG)#(fG)#(mG)#(fU)#(mC)#(fG) | **SS-5** | (fC)#(mU)#(fG)(mG)(fA)(mA)(fA)(mA)(fG)(mC)(fU)(mG)(fA)#(mU)#(fA)-TegChol |
| **D46** | **AS-3** | P(mU)#(fC)#(mC)(fG)(mG)(fU)(mC)(fA)(mC)(fA)(mA)(fC)(mA)#(fU)#(mU)#(fG)#(mU)#(fG)#(mG)#(fU) | **SS-6** | (fA)#(mA)#(fU)(mG)(fU)(mU)(fG)(mU)(fG)(mA)(fC)(mC)(fG)#(mG)#(fA)-TegChol |
| **D47** | **AS-7** | P(**mA**)#(fU)#(mA)(fA)(mU)(fC)(mU)(fC)(mU)(fU)(mU)(fA)(mC)#(fU)#(mG)#(fA)#(mU)#(fA)#(mU)#(fA) | **SS-7** | (fC)#(mA)#(fG)(mU)(fA)(mA)(fA)(mG)(fA)(mG)(fA)(mU)(fU)#(mA)#(**fU**)-TegChol |

**Table S3**. IC50 of each exNA modified siRNAs and potency change

| siRNA# | D1 | D2 | D3 | D4 | D5 |
|---|---|---|---|---|---|
| IC50 | 1.619E-07 | 1.703E-07 | 3.825E-07 | 2.034E-07 | 1.864E-07 |
| IC50 of corresponding control | 7.662E-08 | 3.735E-08 | 7.416E-08 | 7.416E-08 | 7.662E-08 |
| % Potency Change | -111.302532 | -355.957162 | -415.776699 | -174.2718447 | -143.2785174 |
| **siRNA#** | **D6*** | **D7** | **D8*** | **D9** | **D10** |
| IC50 | 8.65E-11 | 2.235E-07 | 1.346E-10 | 0.000000097 | 0.000000111 |
| IC50 of corresponding control | 7.086E-11 | 7.662E-08 | 7.086E-11 | 7.662E-08 | 7.662E-08 |
| % Potency Change | -22.07169066 | -191.6992952 | -89.95201806 | -26.59879927 | -44.87079092 |
| **siRNA#** | **D11** | **D12*** | **D13*** | **D14** | **D15** |
| IC50 | 7.196E-08 | 2.09E-10 | 3.541E-10 | 0.000000175 | 3.119E-07 |
| IC50 of corresponding control | 7.662E-08 | 7.086E-11 | 9.675E-11 | 7.662E-08 | 1.411E-07 |
| % Potency Change | 6.081962934 | -194.9477844 | -265.994832 | -128.3998956 | -121.0489015 |
| **siRNA#** | **D16** | **D17** | **D18*** | **D19** | **D20*** |
| IC50 | 2.782E-07 | 8.911E-08 | 1.924E-10 | 8.779E-08 | 1.589E-10 |
| IC50 of corresponding control | 1.411E-07 | 7.662E-08 | 9.675E-11 | 7.662E-08 | 9.675E-11 |
| % Potency Change | -97.16513111 | -16.30122683 | -98.8630491 | -14.57843905 | -64.2377261 |
| **siRNA#** | **D21** | **D22** | **D23** | **D24** | **D25*** |
| IC50 | 3.365E-08 | 3.713E-08 | 3.884E-08 | 7.226E-08 | 1.192E-10 |
| IC50 of corresponding control | 9.633E-08 | 9.633E-08 | 9.633E-08 | 9.633E-08 | 8.502E-11 |
| % Potency Change | 65.06799543 | 61.45541368 | 59.68026575 | 24.98702377 | -40.20230534 |
| **siRNA#** | **D26** | **D27*** | **D28** | **D29*** | **D30*** |
| IC50 | 1.206E-07 | 8.65E-11 | 1.229E-07 | 1.346E-10 | 2.09E-10 |
| IC50 of corresponding control | 7.416E-08 | 7.019E-11 | 7.662E-08 | 7.019E-11 | 7.019E-11 |
| % Potency Change | -62.62135922 | -23.23692834 | -60.40198382 | -91.76520872 | -197.7632141 |
| **siRNA#** | **D31** | **D32** | **D33** | **D34** | **D35** |
| IC50 | 1.186E-07 | 1.668E-07 | 8.044E-08 | 9.471E-08 | 3.102E-07 |
| IC50 of corresponding control | 1.411E-07 | 7.416E-08 | 7.416E-08 | 7.416E-08 | 2.972E-07 |
| % Potency Change | 15.94613749 | -124.9190939 | -8.468176915 | -27.71035599 | -4.374158816 |
| **siRNA#** | **D36** | **D37** | **D38** | **D39** | |
| IC50 | 4.829E-08 | 4.945E-08 | 2.133E-07 | 3.917E-08 | |
| IC50 of corresponding control | 7.662E-08 | 7.662E-08 | 2.972E-07 | 3.266E-08 | |
| % Potency Change | 36.97468024 | 35.46071522 | 28.23014805 | -19.93263931 | |

**Table S4. Sequence of siRNAs used for mice study**

| Duplex# | Target | Antisense strand (5'->3') | Sense strand (5'->3') |
|---|---|---|---|
| **D48** (NTC) | N/A | VP(mU)#(fA)#(mA)(fU)(mC)(fG)(mU)(fA)(mU)(fU)(mU)(fG)(mU)(fC)(mA)(fA)(mU)#(fC)#(mA)#(ex_mU)#(ex_fU) | (mU)#(fU)#(mG)(fA)(mC)(fA)(mA)(fA)(mU)(fA)(mC)(fG)(mA)(fU)#(mU)#(fA)(dT)(dT)-DCA |
| **D49** | *Htt* | VP(mU)#(fU)#(mA)(fA)(mU)(fC)(mU)(fC)(mU)(fU)(mU)(fA)(mC)(fU)(mG)(fA)(mU)#(fA)#(mU)#(mU)#(fU) | (mU)#(fC)#(mA)(fG)(mU)(fA)(mA)(fA)(mG)(fA)(mG)(fA)(mU)(fU)#(mA)#(fA)(dT)(dT)-DCA |
| **D50** | *Htt* | VP(mU)#(fU)#(mA)(fA)(mU)(fC)(mU)(fC)(mU)(fU)(mU)(fA)(mC)(fU)(mG)(fA)(mU)#(fA)#(mU)#(ex_mU)#(ex_fU) | (mU)#(fC)#(mA)(fG)(mU)(fA)(mA)(fA)(mG)(fA)(mG)(fA)(mU)(fU)#(mA)#(fA)(dT)(dT)-DCA |
| **D51** (NTC) | N/A | VP(mU)#(fA)#(mA)(fU)(fC)(fG)(mU)(fA)(mU)(fU)(mU)(fG)(mU)(fC)(mA)(fA)(mU)(mC)(mA)#(mU)#(fU) | (mU)#(mU)#(mG)(fA)(mC)(fA)(mA)(fA)(mU)(fA)(mC)(mG)(mA)(fU)#(mU)#(mA)(dT)(dT)-DCA |
| **D52** | *Mstn* | VP(mU)#(fU)#(mA)(fU)(fU)(fA)(mU)(fU)(mU)(fG)(mU)(fU)(mC)(fU)#(mU)#(fU)#(mG)#(mC)#(mC)#(mU)#(fU) | (mA)#(mA)#(mA)(fG)(mA)(fA)(mC)(fA)(mA)(fA)(mU)(mA)(mA)(fU)#(mA)#(mA)(dT)(dT)-DCA |
| **D53** | *Mstn* | VP(mU)#(fU)#(mA)(fU)(fU)(fA)(mU)(fU)(mU)(fG)(mU)(fU)(mC)(fU)#(mU)#(fU)#(mG)#(mC)#(mC)#(ex-mU)#(ex-fU) | (mA)#(mA)#(mA)(fG)(mA)(fA)(mC)(fA)(mA)(fA)(mU)(mA)(mA)(fU)#(mA)#(mA)(dT)(dT)-DCA |
| **D54** (NTC) | N/A | VP(mU)#(fA)#(mA)(mU)(mC)(fG)(mU)(mA)(mU)(mU)(mU)(mG)(mU)(fC)#(mA)#(fA)#(mU)#(mC)#(mA)#(fU)#(mA) | (mU)#(mU)#(mG)(mA)(mC)(fA)(fA)(fA)(mU)(fA)(mC)(mG)(mA)(mU)#(mU)#(mA)-DIO |
| **D55** | *ApoE* | VP(mU)#(fU)#(mG)(mG)(mA)(fU)(mA)(mU)(mG)(mG)(mA)(mU)(mG)(fU)(mU)(fG)(mU)(mU)(mU)#(mU)#(mU) | (mC)#(mA)#(mA)(mC)(mA)(mU)(mC)(fC)(mA)(fU)(fA)(fU)(mC)(mC)#(mA)#(mA)-DIO |
| **D56** | *ApoE* | VP(mU)#(fU)#(mG)(mG)(mA)(fU)(mA)(mU)(mG)(mG)(mA)(mU)(mG)(fU)(mU)(fG)(mU)(mU)(mU)#(ex-mU)#(ex-mU) | (mC)#(mA)#(mA)(mC)(mA)(mU)(mC)(fC)(mA)(fU)(fA)(fU)(mC)(mC)#(mA)#(mA)-DIO |
| **D57** (NTC) | N/A | VP(mU)#(fA)#(mA)(fU)(fC)(fG)(mU)(fA)(mU)(fU)(mU)(fG)(mU)(fC)(mA)(fA)(mU)(mC)(mA)#(mU)#(fU) | (mU)#(mU)#(mG)(fA)(mC)(mA)(mA)(fA)(mU)(fA)(mC)(fG)(mA)(fU)#(mU)#(mA)-DIO |
| **D58** | *Htt* | VP(mU)#(fU)#(mA)(fA)(fU)(fC)(mU)(fC)(mU)(fU)(mU)(fA)(mC)(fU)(mG)(fA)(mU)(mA)(mU)#(mU)#(fU) | (mU)#(mC)#(mA)(fG)(mU)(mA)(mA)(fA)(mG)(fA)(mG)(fA)(mU)(fU)#(mA)#(mA)-DIO |
| **D59** | *Htt* | VP(mU)#(fU)#(mA)(fA)(fU)(fC)(mU)(fC)(mU)(fU)(mU)(fA)(mC)(fU)(mG)(fA)(mU)(mA)(mU)#(ex-mU)#(ex-fU) | (mU)#(mC)#(mA)(fG)(mU)(mA)(mA)(fA)(mG)(fA)(mG)(fA)(mU)(fU)#(mA)#(mA)-DIO |

Chemical modifications are designated as follows, "#": Phosphorothioate bond, "m": 2'-O-Methyl, "f": 2'-Fluoro, "VP": 5'-(*E*)-Vinylphosphonate, DCA: Docosanoic acid conjugate, "DIO": Divalent-siRNA. **D48**, **D51**, **D54**, **D57** are NTC for **D49/50**, **D52/53**, **D55/56**, **D58/59**, respectively.

**Table S5.** The sequence of oligonucleotides used for thermal melting analysis

| ON # | Sequence (5′ -> 3′)[a] | Calcd mass $[M-H]^-$ | Found mass |
|---|---|---|---|
| **ON1** | (rC)(rG)(rC)(rA)(rU)(rU)(rA)(rG)(rC)(rC)(rG) | 3465.1 | 3467.0 |
| **ON2** | (rC)(rG)(rC)(rA)(rU)(rxU)(rA)(rG)(rC)(rC)(rG) | 3479.6 | 3478.5 |
| **cRNA1** | (rC)(rG)(rG)(rC)(rU)(rA)(rA)(rU)(rG)(rC)(rG) | 3505.2 | 3504.5 |
| **cRNA2** | (rC)(rG)(rG)(rC)(rU)(rG)(rA)(rU)(rG)(rC)(rG) | 3481.1 | 3480.5 |
| **cRNA3** | (rC)(rG)(rG)(rC)(rU)(rU)(rA)(rU)(rG)(rC)(rG) | 3482.1 | 3481.5 |
| **cRNA4** | (rC)(rG)(rG)(rC)(rU)(rC)(rA)(rU)(rG)(rC)(rG) | 3521.2 | 3520.4 |

[a] (rN) : 2'-ribonucleosides, (ex-rU): exNA-uridine (2'-OH).

**Table S6**. Oligonucleotide used for 3'-exonuclease resistance assay

| ON# | Sequence (5'->3') | Calcd mass [M-H]$^-$ | Found mass |
|---|---|---|---|
| **ON3** | (dT)$_{18}$(mU)(mU) | 6053.9 | 6053.0 |
| **ON4** | (dT)$_{18}$(**ex-mU**)(**ex-mU**) | 6113.9 | 6113.0 |
| **ON5** | (dT)$_{18}$ **#**(mU)**#**(mU) | 6086.1 | 6085.9 |
| **ON6** | (dT)$_{18}$ **#**(**ex-mU**)**#**(**ex-mU**) | 6114.1 | 6114.0 |

**Table S7**. Oligonucleotide used for 5'-exonuclease resistance assay

| ON# | Sequence (5'->3') | Calcd mass [M-H]$^-$ | Found mass |
|---|---|---|---|
| **ON7** | (mU)(dT)(dT)$_{18}$ | 6037.9 | 6036.9 |
| **ON8** | (mU)#(dT)(dT)$_{18}$ | 6054.0 | 6053.9 |
| **ON9** | (**ex-mU**)(dT)(dT)$_{18}$ | 6052.0 | 6050.9 |
| **ON10** | (mU)(mU)(dT)$_{18}$ | 6053.9 | 6053.9 |
| **ON11** | (mU)#(mU)(dT)$_{18}$ | 6070.0 | 6070.0 |
| **ON12** | (mU)(**ex-mU**)(dT)$_{18}$ | 6067.9 | 6067.9 |

**Table S8. Mass analysis of exNA modified antisense strands used in exNA walk study**

| Name | Sequence (5'->3') | Mass calcd [M-H]$^-$ | Mass found |
|---|---|---|---|
| ex-AS1 | P(**ex-mU**)#(fU)#(mA)(fA)(mU)(fC)(mU)(fC)(mU)(fU)(mU)(fA)(mC)#(fU)#(mG)#(fA)#(mU)#(fA)#(mU)#(fA) | 6633.4 | 6633.7 |
| ex-AS2 | P(mU)#(**ex-fU**)#(mA)(fA)(mU)(fC)(mU)(fC)(mU)(fU)(mU)(fA)(mC)#(fU)#(mG)#(fA)#(mU)#(fA)#(mU)#(fA) | 6633.4 | 6633.7 |
| ex-AS3 | P(mU)#(fU)#(**ex-mU**)(fU)(mU)(fA)(mA)(fA)(mU)(fC)(mC)(fU)(mG)#(fA)#(mG)#(fA)#(mA)#(fG)#(mA)#(fA) | 6758.6 | 6758.8 |
| ex-AS4 | P(mU)#(fU)#(mU)(**ex-fU**)(mU)(fA)(mA)(fA)(mU)(fC)(mC)(fU)(mG)#(fA)#(mG)#(fA)#(mA)#(fG)#(mA)#(fA) | 6758.6 | 6758.8 |
| ex-AS5 | P(mU)#(fU)#(mA)(fA)(**ex-mU**)(fC)(mU)(fC)(mU)(fU)(mU)(fA)(mC)#(fU)#(mG)#(fA)#(mU)#(fA)#(mU)#(fA) | 6633.4 | 6633.7 |
| ex-AS6 | P(mU)#(fC)#(mC)(fA)(mC)(**ex-fU**)(mA)(fU)(mG)(fU)(mU)(fU)(mU)#(fC)#(mA)#(fC)#(mA)#(fU)#(mA)#(fU) | 6608.4 | 6608.7 |
| ex-AS7 | P(mU)#(fU)#(mA)(fA)(mU)(fC)(**ex-mU**)(fC)(mU)(fU)(mU)(fA)(mC)#(fU)#(mG)#(fA)#(mU)#(fA)#(mU)#(fA) | 6633.4 | 6632.7 |
| ex-AS8 | P(mU)#(fC)#(mC)(fA)(mC)(fU)(mA)(**ex-fU**)(mG)(fU)(mU)(fU)(mU)#(fC)#(mA)#(fC)#(mA)#(fU)#(mA)#(fU) | 6608.4 | 6608.7 |
| ex-AS9 | P(mU)#(fU)#(mA)(fA)(mU)(fC)(mU)(fC)(**ex-mU**)(fU)(mU)(fA)(mC)#(fU)#(mG)#(fA)#(mU)#(fA)#(mU)#(fA) | 6633.4 | 6633.7 |
| ex-AS10 | P(mU)#(fU)#(mA)(fA)(mU)(fC)(mU)(fC)(mU)(**ex-fU**)(mU)(fA)(mC)#(fU)#(mG)#(fA)#(mU)#(fA)#(mU)#(fA) | 6633.4 | 6633.7 |
| ex-AS11 | P(mU)#(fU)#(mA)(fA)(mU)(fC)(mU)(fC)(mU)(fU)(**ex-mU**)(fA)(mC)#(fU)#(mG)#(fA)#(mU)#(fA)#(mU)#(fA) | 6633.4 | 663.7 |
| ex-AS12 | P(mU)#(fC)#(mC)(fA)(mC)(fU)(mA)(fU)(mG)(fU)(mU)(**ex-fU**)(mU)#(fC)#(mA)#(fC)#(mA)#(fU)#(mA)#(fU) | 6608.4 | 6608.7 |
| ex-AS13 | P(mU)#(fC)#(mC)(fA)(mC)(fU)(mA)(fU)(mG)(fU)(mU)(fU)(**ex-mU**)#(fC)#(mA)#(fC)#(mA)#(fU)#(mA)#(fU) | 6608.4 | 6608.7 |
| ex-AS14 | P(mU)#(fU)#(mA)(fA)(mU)(fC)(mU)(fC)(mU)(fU)(mU)(fA)(mC)#(**ex-fU**)#(mG)#(fA)#(mU)#(fA)#(mU)#(fA) | 6633.4 | 6633.7 |

| | | | |
|---|---|---|---|
| ex-AS15 | P(mU)#(fG)#(mC)(fC)(mU)(fA)(mA)(fG)(mA)(fG)(mC)(fA)(mC)#(fA)#(**ex-mU**)#(fU)#(mU)#(fA)#(mG)#(fU) | 6749.6 | 6749.8 |
| ex-AS16 | P(mU)#(fG)#(mC)(fC)(mU)(fA)(mA)(fG)(mA)(fG)(mC)(fA)(mC)#(fA)#(mU)#(**ex-fU**)#(mU)#(fA)#(mG)#(fU) | 6749.6 | 6749.8 |
| ex-AS17 | P(mU)#(fU)#(mA)(fA)(mU)(fC)(mU)(fC)(mU)(fU)(mU)(fA)(mC)#(fU)#(mG)#(fA)#(**ex-mU**)#(fA)#(mU)#(fA) | 6633.4 | 6633.7 |
| ex-AS18 | P(mU)#(fC)#(mC)(fA)(mC)(fU)(mA)(fU)(mG)(fU)(mU)(fU)(mU)#(fC)#(mA)#(fC)#(mA)#(**ex-fU**)#(mA)#(fU) | 6608.4 | 6608.7 |
| ex-AS19 | P(mU)#(fU)#(mA)(fA)(mU)(fC)(mU)(fC)(mU)(fU)(mU)(fA)(mC)#(fU)#(mG)#(fA)#(mU)#(fA)#(**ex-mU**)#(fA) | 6633.4 | 6633.7 |
| ex-AS20 | P(mU)#(fC)#(mC)(fA)(mC)(fU)(mA)(fU)(mG)(fU)(mU)(fU)(mU)#(fC)#(mA)#(fC)#(mA)#(fU)#(mA)#(**ex-fU**) | 6608.4 | 6608.7 |
| ex-AS21 | P(mU)#(fU)#(mA)(fA)(mU)(fC)(mU)(fC)(mU)(fU)(mU)(fA)(mC)#(fU)#(mG)#(fA)#(mU)#(fU)#(mU)#(**ex-fU**) | 6587.4 | 6587.6 |
| ex-AS22 | P(mU)#(fU)#(mA)(fA)(mU)(fC)(mU)(fC)(mU)(fU)(mU)(fA)(mC)#(fU)#(mG)#(fA)#(mU)#(fU)#(**ex-mU**)<br>#(**ex-fU**) | 6601.4 | 6601.6 |
| ex-AS23 | P(mU)#(fU)#(mA)(fA)(mU)(fC)(mU)(fC)(mU)(fU)(mU)(fA)(mC)#(fU)#(mG)#(fA)#(mU)#(**ex-fU**)<br>#(**ex-mU**)#(**ex-fU**) | 6615.4 | 6615.7 |
| ex-AS24 | P(mU)#(fU)#(mA)(fA)(mU)(fC)(mU)(fC)(mU)(fU)(mU)(fA)(mC)#(fU)#(mG)#(fA)#(**ex-mU**)#(**ex-fU**)<br>#(**ex-mU**)#(**ex-fU**) | 6629.4 | 6629.7 |

P: 5'-phosphate, "m": 2'-OMe, "f": 2'-fluoro, "ex-mU": 2'-OMe-exNA-uridine, "ex-fU": 2'-F-exNA-uridine, #: phosphorothioate

**Table S9.** **Mass analysis of exNA modified sense strands used in exNA walk study**

| Name | **Sequence (Sense strands)** | Mass calcd [M-H]$^-$ | Mass found |
|---|---|---|---|
| ex-SS1 | (**ex-fU**)#(mG)#(fA)(mA)(fA)(mA)(fC)(mA)(fU)(mA)(fG)(mU)(fG)#(mG)#(fA)-TegChol | 5795.30 | 5795.2 |
| ex-SS2 | (fC)#(**ex-mU**)#(fC)(mA)(fG)(mG)(fA)(mU)(fU)(mU)(fA)(mA)(fA)#(mA)#(fA)-TegChol | 5716.30 | 5716.2 |
| ex-SS3 | (fA)#(mA)#(**ex-fU**)(mG)(fU)(mU)(fG)(mU)(fG)(mA)(fC)(mC)(fG)#(mG)#(fA)-TegChol | 5764.30 | 5764.1 |
| ex-SS4 | (fC)#(mA)#(fG)(**ex-mU**)(fA)(mA)(fA)(mG)(fA)(mG)(fA)(mU)(fU)#(mA)#(fA)-TegChol | 5779.40 | 5778.2 |
| ex-SS5 | (fA)#(mA)#(fU)(mG)(**ex-fU**)(mU)(fG)(mU)(fG)(mA)(fC)(mC)(fG)#(mG)#(fA)-TegChol | 5764.30 | 5763.2 |
| ex-SS6 | (fA)#(mA)#(fU)(mG)(fU)(**ex-mU**)(fG)(mU)(fG)(mA)(fC)(mC)(fG)#(mG)#(fA)-TegChol | 5764.30 | 5764.2 |
| ex-SS7 | (fA)#(mU)#(fG)(mU)(fG)(mC)(**ex-fU**)(mC)(fU)(mU)(fA)(mG)(fG)#(mC)#(fA)-TegChol | 5701.20 | 5701.1 |
| ex-SS8 | (fC)#(mU)#(fC)(mA)(fG)(mG)(fA)(**ex-mU**)(fU)(mU)(fA)(mA)(fA)#(mA)#(fA)-TegChol | 5716.30 | 5716.2 |
| ex-SS9 | (fC)#(mU)#(fC)(mA)(fG)(mG)(fA)(mU)(**ex-fU**)(mU)(fA)(mA)(fA)#(mA)#(fA)-TegChol | 5716.30 | 5716.2 |
| ex-SS10 | (fC)#(mU)#(fC)(mA)(fG)(mG)(fA)(mU)(fU)(**ex-mU**)(fA)(mA)(fA)#(mA)#(fA)-TegChol | 5716.30 | 5716.1 |
| ex-SS11 | (fC)#(mU)#(fG)(mG)(fA)(mA)(fA)(mA)(fG)(mC)(**ex-fU**)(mG)(fA)#(mU)#(fA)-TegChol | 5771.30 | 5771.2 |
| ex-SS12 | (fC)#(mA)#(fG)(mU)(fA)(mA)(fA)(mG)(fA)(mG)(fA)(**ex-mU**)(fU)#(mA)#(fA)-TegChol | 5779.40 | 5779.2 |
| ex-SS13 | (fC)#(mA)#(fG)(mU)(fA)(mA)(fA)(mG)(fA)(mG)(fA)(mU)(**ex-fU**)#(mA)#(fA)-TegChol | 5779.30 | 5779.2 |
| ex-SS14 | (fC)#(mU)#(fG)(mG)(fA)(mA)(fA)(mA)(fG)(mC)(fU)(mG)(fA)#(**ex-mU**)#(fA)-TegChol | 5771.30 | 5771.2 |
| ex-SS15 | (fC)#(mA)#(fG)(mU)(fA)(mA)(fA)(mG)(fA)(mG)(fA)(mU)(fU)#(mA)#(**ex-fU**)-TegChol | 5756.30 | 5756.2 |

**Table S10. Mass analysis of control antisense and sense strands used in exNA walk study**

| Name | Sequence (Antisense strands) | Mass calcd [M-H]$^-$ | Mass found |
|---|---|---|---|
| **AS-1** | P(mU)#(fC)#(mC)(fA)(mC)(fU)(mA)(fU)(mG)(fU)(mU)(fU)(mU)#(fC)#(mA)#(fC)#(mA)#(fU)#(mA)#(fU) | 6595.40 | 6594.7 |
| **AS-2** | P(mU)#(fU)#(mU)(fU)(mU)(fA)(mA)(fA)(mU)(fC)(mC)(fU)(mG)#(fA)#(mG)#(fA)#(mA)#(fG)#(mA)#(fA) | 6745.60 | 6744.7 |
| **AS-3** | P(mU)#(fC)#(mC)(fG)(mG)(fU)(mC)(fA)(mC)(fA)(mA)(fC)(mA)#(fU)#(mU)#(fG)#(mU)#(fG)#(mG)#(fU) | 6728.60 | 6727.7 |
| **AS-4** | P(mU)#(fU)#(mA)(fA)(mU)(fC)(mU)(fC)(mU)(fU)(mU)(fA)(mC)#(fU)#(mG)#(fA)#(mU)#(fA)#(mU)#(fA) | 6620.40 | 6619.7 |
| **AS-5** | P(mU)#(fG)#(mC)(fC)(mU)(fA)(mA)(fG)(mA)(fG)(mC)(fA)(mC)#(fA)#(mU)#(fU)#(mU)#(fA)#(mG)#(fU) | 6736.40 | 6735.7 |
| **AS-6** | P(mU)#(fA)#(mU)(fC)(mA)(fG)(mC)(fU)(mU)(fU)(mU)(fC)(mC)#(fA)#(mG)#(fG)#(mG)#(fU)#(mC)#(fG) | 6705.60 | 6704.7 |
| **AS-7** | P(mA)#(fU)#(mA)(fA)(mU)(fC)(mU)(fC)(mU)(fU)(mU)(fA)(mC)#(fU)#(mG)#(fA)#(mU)#(fA)#(mU)#(fA) | 6643.50 | 6642.7 |
| **AS-8** | P(mU)#(fU)#(mA)(fA)(mU)(fC)(mU)(fC)(mU)(fU)(mU)(fA)(mC)#(fU)#(mG)#(fA)#(mU)#(fU)#(mU)#(fU) | 6574.40 | 6573.6 |
| **SS-1** | (fC)#(mA)#(fG)(mU)(fA)(mA)(fA)(mG)(fA)(mG)(fA)(mU)(fU)#(mA)#(fA)-TegChol | 5765.30 | 5765.2 |
| **SS-2** | (fC)#(mU)#(fC)(mA)(fG)(mG)(fA)(mU)(fU)(mU)(fA)(mA)(fA)#(mA)#(fA)-TegChol | 5702.30 | 5702.2 |
| **SS-4** | (fU)#(mG)#(fA)(mA)(fA)(mA)(fC)(mA)(fU)(mA)(fG)(mU)(fG)#(mG)#(fA)-TegChol | 5781.30 | 5781.2 |
| **SS-5** | (fA)#(mU)#(fG)(mU)(fG)(mC)(fU)(mC)(fU)(mU)(fA)(mG)(fG)#(mC)#(fA)-TegChol | 5687.20 | 5686.1 |
| **SS-6** | (fC)#(mU)#(fG)(mG)(fA)(mA)(fA)(mA)(fG)(mC)(fU)(mG)(fA)#(mU)#(fA)-TegChol | 5757.30 | 5757.2 |
| **SS-7** | (fA)#(mA)#(fU)(mG)(fU)(mU)(fG)(mU)(fG)(mA)(fC)(mC)(fG)#(mG)#(fA)-TegChol | 5750.30 | 5750.1 |
| **SS-8** | (fC)#(mA)#(fG)(mU)(fA)(mA)(fA)(mG)(fA)(mG)(fA)(mU)(fU)#(mA)#(fU)-TegChol | 5742.30 | 5742.2 |

**Table S11. Mass analysis of oligonucleotides used for in vivo study**

| Name | Sequence (5'->3') | Calcd mass [M-H]$^-$ | Found mass |
|---|---|---|---|
| D48-AS | VP(mU)#(fU)#(mA)(fA)(mU)(fC)(mU)(fC)(mU)(fU)(mU)(fA)(mC)(fU)(mG)(fA)(mU)#(fA)#(mU)#(mU)#(fU) | 6864.4 | 6864.8 |
| D49-AS | VP(mU)#(fU)#(mA)(fA)(mU)(fC)(mU)(fC)(mU)(fU)(mU)(fA)(mC)(fU)(mG)(fA)(mU)#(fA)#(mU)#(ex-mU)#(ex-fU) | 6892.4 | 6892.8 |
| D50-AS (NTC) | VP(mU)#(fA)#(mA)(fU)(mC)(fG)(mU)(fA)(mU)(fU)(mU)(fG)(mU)(fC)(mA)(fA)(mU)#(fC)#(mA)#(ex-mU)#(ex-fU) | 6954.5 | 6954.9 |
| D55-AS | VP(mU)#(fU)#(mG)(mG)(mA)(fU)(mA)(mU)(mG)(mG)(mA)(mU)(mG)(fU)(mU)(fG)(mU)(mU)(mU)#(mU)#(mU) | 7056.6 | 7056.9 |
| D56-AS | VP(mU)#(fU)#(mG)(mG)(mA)(fU)(mA)(mU)(mG)(mG)(mA)(mU)(mG)(fU)(mU)(fG)(mU)(mU)(mU)#(ex-mU)#(ex-mU) | 7084.6 | 7084.9 |
| *ApoE*-di-sense strand | (mC)#(mA)#(mA)(mC)(mA)(mU)(mC)(fC)(mA)(fU)(fA)(fU)(mC)(mC)#(mA)#(mA)-Divalent | 10857.3 | 10857.6 |
| D58-AS | VP(mU)#(fU)#(mA)(fA)(fU)(fC)(mU)(fC)(mU)(fU)(mU)(fA)(mC)(fU)(mG)(fA)(mU)(mA)(mU)#(mU)#(fU) | 6832.2 | 6832.8 |
| D59-AS | VP(mU)#(fU)#(mA)(fA)(fU)(fC)(mU)(fC)(mU)(fU)(mU)(fA)(mC)(fU)(mG)(fA)(mU)(mA)(mU)#(ex-mU)#(ex-fU) | 6860.3 | 6860.9 |
| *Htt*-di-sense strand | (mU)#(mC)#(mA)(fG)(mU)(mA)(mA)(fA)(mG)(fA)(mG)(fA)(mU)(fU)#(mA)#(mA)-Divalent | 11123.4 | 11123.8 |

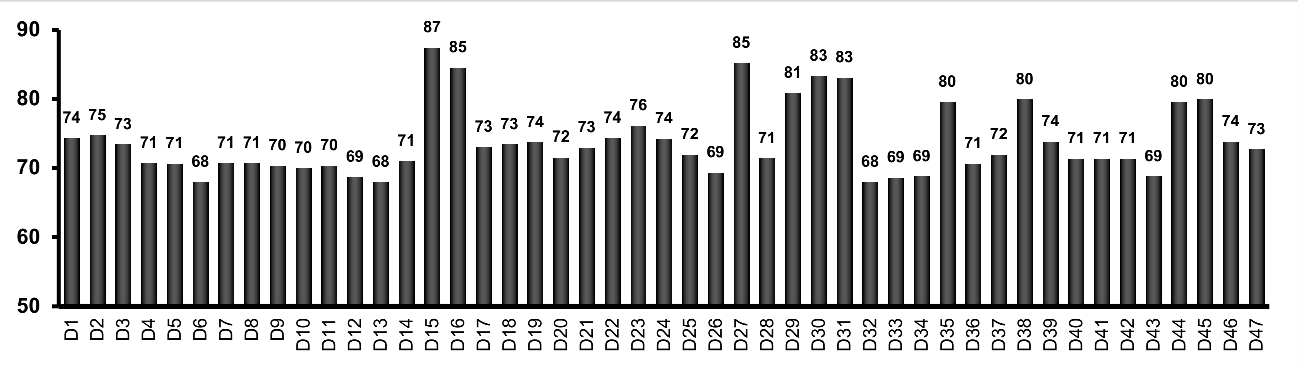


**Fig. S1** *T*m Values of each siRNA duplexes (**D1-47**) used in in-vitro exNA walk. Melting temperature analysis was conducted in a buffer 10mM Sodium phosphate buffer (pH 7.2) containing 100 mM NaCl, 0.1 mM EDTA and 1 mM guide/sense strand. $T_m$ values were average values determined in triplicate experiments.

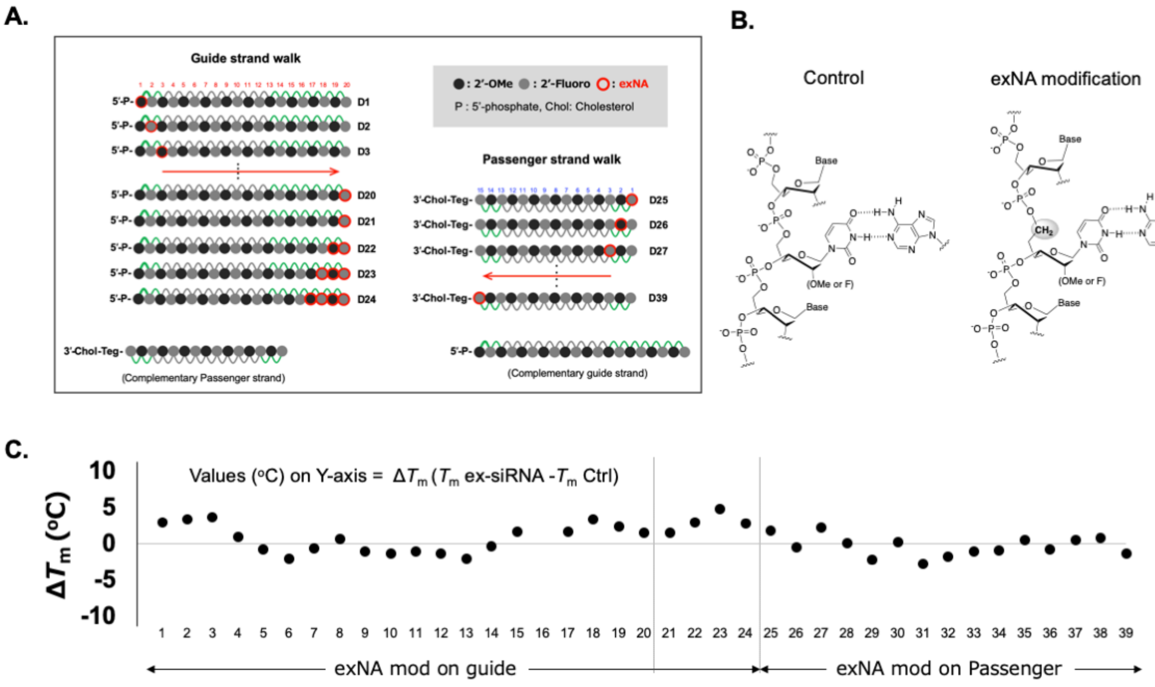


**Fig. S2 Delta-*Tm* of each siRNA duplexes.** (A) Schematic of exNA walk and corresponding Duplex numbers. (B) Structural difference between duplex with canonical backbone and with exNA backbone. (C) delta-Tm values for each exNA modification positions on guide and passenger strand. Calculation of delta-*Tm* was conducted based on *Tm* values shown in **Fig. S1**.

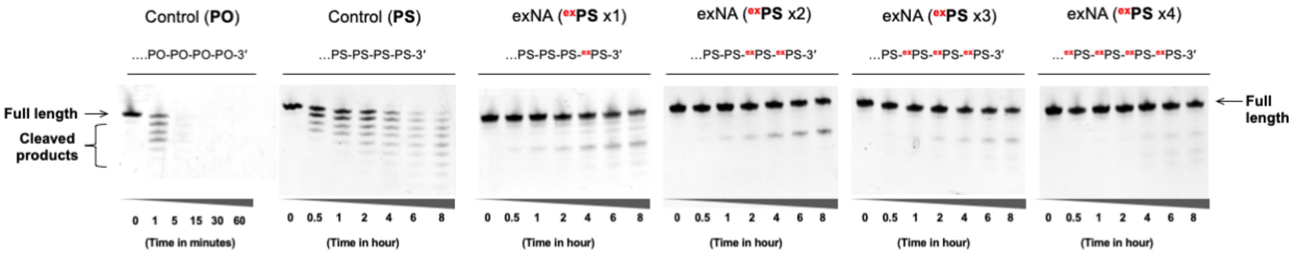


| | |
|---|---|
| Control (PO) | P(mU)(fU)(mA)(fA)(mU)(fC)(mU)(fC)(mU)(fU)(mU)(fA)(mC)(fU)(mG)(fA)(mU)(fU)(mU)(fU) |
| Control (PS) | P(mU)#(fU)#(mA)(fA)(mU)(fC)(mU)(fC)(mU)(fU)(mU)(fA)(mC)#(fU)#(mG)#(fA)#(mU)#(fU)#(mU)#(fU) |
| exNA (exPS x1) | P(mU)#(fU)#(mA)(fA)(mU)(fC)(mU)(fC)(mU)(fU)(mU)(fA)(mC)#(fU)#(mG)#(fA)#(mU)#(fU)#(mU)#(ex_fU) |
| exNA (exPS x2) | P(mU)#(fU)#(mA)(fA)(mU)(fC)(mU)(fC)(mU)(fU)(mU)(fA)(mC)#(fU)#(mG)#(fA)#(mU)#(fU)#(ex_mU)#(ex_fU) |
| exNA (exPS x3) | P(mU)#(fU)#(mA)(fA)(mU)(fC)(mU)(fC)(mU)(fU)(mU)(fA)(mC)#(fU)#(mG)#(fA)#(mU)#(ex_fU)#(ex_mU)#(ex_fU) |
| exNA (exPS x4) | P(mU)#(fU)#(mA)(fA)(mU)(fC)(mU)(fC)(mU)(fU)(mU)(fA)(mC)#(fU)#(mG)#(fA)#(ex_mU)#(ex_fU)#(ex_mU)#(ex_fU) |

Chemical modifications are designated as follows, "#" –phosphorothioate bond, "m": 2'-O-Methyl, "f": 2'-Fluoro, "P": 5' Phosphate, "ex_": exNA

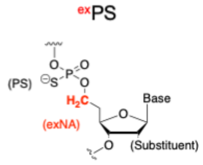


**Fig. S3** 17.5 uM oligonucleotide was incubated in a 10 mM Tris-HCl (pH 8) buffer containing 2 mM $MgCl_2$ and 20 mU/mL Snake Venom Phosphodiesterase I. Aliquot at each time point was dissolved into 95% formamide containing 5% EDTA

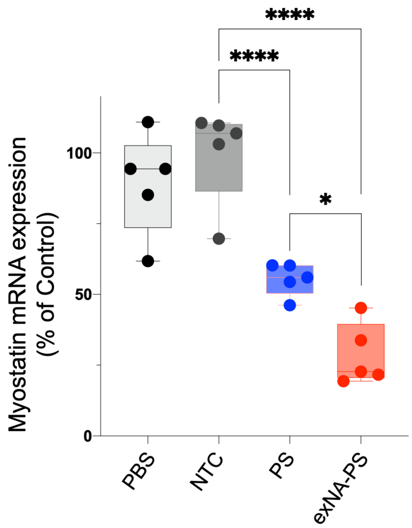


**Fig. S4 exNA-PS modification enhances Myostatin-targeting siRNA efficacy.** DCA conjugated siRNAs (D51, 52, and 53) were administered to FVB mice (female, 7-8 weeks old) at 20mg/kg dosage twice (3 days apart, s.c.). Mice was sacrificed at 1 week post injection, and muscle quadricep was collected. Myostatin mRNA level was quantified by QuantiGene. NTC: Non-targeting control (**D51**), PS: non-exNA PS modified siRNA (**D52**), exNA-PS: exNA-PS modified siRNA (**D53**). For sequences, see **Table S9**.
